# Assembling membraneless organelles from *de novo* designed proteins

**DOI:** 10.1101/2023.04.18.537322

**Authors:** Alexander T. Hilditch, Andrey Romanyuk, Stephen J. Cross, Richard Obexer, Jennifer J. McManus, Derek N. Woolfson

## Abstract

Recent advances in *de novo* protein design have delivered a diversity of discrete *de novo* protein structures and complexes. A new challenge for the field is to use these designs directly in cells to intervene in biological process and augment natural systems. The bottom-up design of self-assembled objects like microcompartments and membraneless organelles is one such challenge, which also presents opportunities for chemical and synthetic biology. Here, we describe the design of genetically encoded polypeptides that form membraneless organelles in *Escherichia coli* (*E. coli*). To do this, we combine *de novo* α-helical sequences, intrinsically disordered linkers, and client proteins in single-polypeptide constructs. We tailor the properties of the helical regions to shift protein assembly from diffusion-limited assemblies to dynamic condensates. The designs are characterised in cells and *in vitro* using biophysical and soft-matter physics methods. Finally, we use the designed polypeptide to co-compartmentalise a functional enzyme pair in *E. coli*.

## INTRODUCTION

Biomolecular condensates are an emerging paradigm for a collective type of biomacromolecular organisation in cells. The presence of dynamic cellular compartments known as membraneless organelles (MLOs) has been known for some time.^1,2^ However, the widespread occurrence and utility of the phenomenon in biological systems, and specifically within cells have only recently become apparent.^3,4^ Biomolecular condensates can take diverse forms including: amorphous aggregates, viscous liquids and gels, liquid-liquid phase-separated (LLPS) compartments, and complex coacervates formed by protein-nucleic acid interactions.^5,6^ Each mode of condensation has different physical properties, and therefore the specific organisation of macromolecules within the condensate has functional consequences.^7^ LLPS is of particular interest because it can lead to highly dynamic and reversible cellular compartments that can respond to internal or external stimuli.^8^ LLPS occurs when soluble macromolecules reversibly separate into de-mixed liquid phases, leaving one richer in the macromolecules than the other (Fig. 1a).^9^ LLPS creates dense macromolecular phases that can accommodate diverse clients at high local concentrations, while permitting small molecules, proteins, and nucleic acids to diffuse between the organelle and its surroundings.^10^

**Figure 1:**
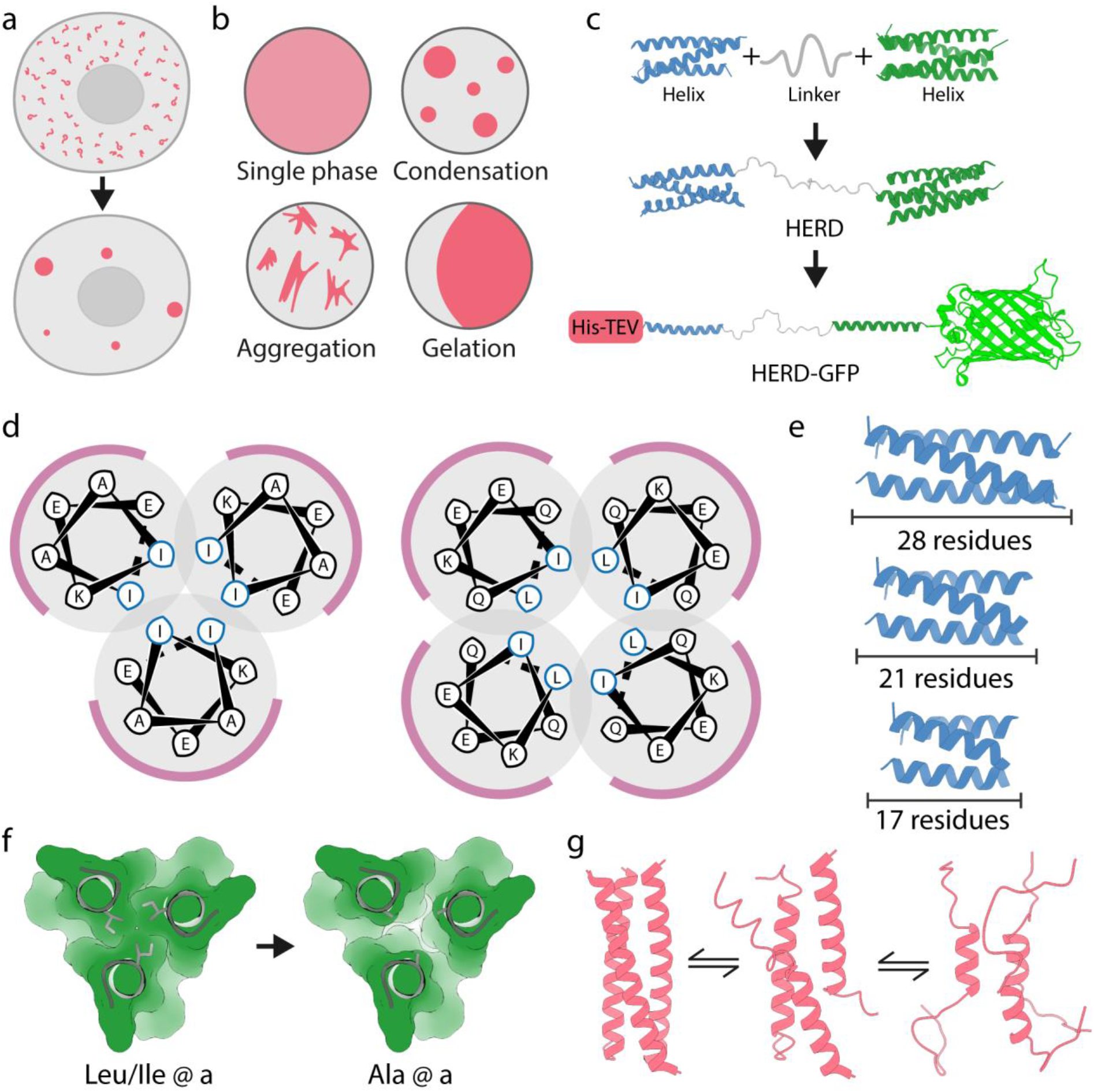
Design and assembly of *de novo* polypeptides for biomolecular condensation. **a,** Cartoon for membraneless-organelle formation in cells; *i.e.*, protein condensation leading to the formation of de-mixed droplets. **b,** Protein solutions can form a single phase, or phase-separated systems including condensates, aggregates, and gels. **c,** HERD design strategy for phase separation by concatenation of *de novo* CCs. **d,** Helical wheels of the heptad (7-residue) repeats for trimeric (left) and tetrameric (right) CCs with hydrophobic interface residues in blue and solvent exposed residues in white. **e – g,** Weakening PPIs by: **e**, truncating the helical CC lengths; **f**, disrupting packing in the hydrophobic core through Ile/Leu (left) to Ala (right) mutations; and **g**, reducing helical propensity by replacing surface residues to those with a low helical propensities.

The ubiquity and utility of LLPS and MLOs in biology has brought the phenomenon to the attention of synthetic biologists.^11^ Their aim is clear: to build artificial phase-separated compartments within cells to provide new and engineerable routes to functional MLOs. Indeed, artificially induced protein condensation has been demonstrated by exploiting the properties of natural and engineered intrinsically disordered proteins (IDPs).^12-18^ As an alternative to using natural sequences, here we advocate bottom-up *de novo* design of polypeptides to promote protein condensation in cells. This uses weak polypeptide-polypeptide interactions (PPIs) to drive condensation. It presents a programmable platform orthogonal to the host proteome, with the potential to expand the capabilities of LLPS and MLOs in synthetic biology. With the coming of age of *de novo* protein design,^19^ researchers are now exploring the construction of protein assemblies that interface with and augment biology.^20^ These include small self-assembled polypeptide-based objects (origamis);^21^ fibrous materials for organising proteins and reporting on cellular events;^22-24^ and the rational and computational design of large peptide- and protein-based cages for cell delivery.^25^ The design of peptides or proteins for LLPS would explore unchartered design space by exploiting weak and structurally less-well-defined interactions. *De novo* proteins, such as our own set of *de novo* α-helical coiled coils (CCs),^26,27^ are good starting points for creating new assemblies^28^ due to their defined interactions and orthogonality to natural proteomes.^29^ CCs provide high valencies encoded in short helical sequences. Therefore, they have the potential to mimic the high-valency interactions of natural proteins that undergo LLPS.^30^ Further, our understanding of sequence-to-structure relationships for CCs presents a tractable route towards engineering the PPIs that they make and, thus, their collective solution behaviour.^31,32^

Here, we present the *de novo* design and characterisation of genetically encoded polypeptides that form dynamic droplets under physiological conditions in *E. coli*. We describe the concatenation of multivalent *de novo* CCs. The properties of these helical regions are then tuned to direct weakened PPIs leading to condensation consistent with LLPS. Droplet formation is reversible with an upper critical solution temperature (UCST). The PPIs are weakly attractive with an interaction parameter, *k*_*D*_, consistent with natural proteins that undergo LLPS. Interestingly, LLPS is triggerable within a physiologically accessible temperature range, which we exploit to modulate the material properties of droplets directly within bacteria. Finally, we demonstrate the potential of our *de novo* polypeptide system to generate functional organelle-like compartments in *E. coli* by co-compartmentalising different client proteins including two enzymes to produce indigo in the host cells.

## Results and discussion

### *De novo* design delivers subcellular protein condensates

To generate a modular polypeptide that promotes biomolecular condensation in cells, we focused on emulating high-valency protein-protein interactions of the sticker-spacer paradigm for natural condensates and hydrogels.^33,34^ To do this, we concatenated two α-helical CCs via a flexible linker to create helical-repeat domains (HERD; Fig. 1b,c). Specifically, we used extant *de novo* trimeric (CC-Tri)^26^ and tetrameric (CC-Tet)^35^ CCs as the stickers (helical repeats HR1 and HR2), and a flexible 25-residue linker as the spacer. The linker design was guided by amino-acid propensities observed in natural intrinsically disordered proteins (IDPs),^36^ and we targeted a net-zero charge and high hydrophilicity. Glycine residues were used as helical caps to prevent helical readthrough into the linker (Supplementary Fig. 1).^37^ The overall pI of the HERD was lowered from 9.4 to 4.7 by exchanging lysine (Lys) residues for glutamate (Glu) in the HRs to avoid interactions with intracellular nucleic acids and potential coacervate formation (Fig. 1d).^38,39^ As an initial client protein and to facilitate imaging, the monomeric fluorescent protein mEmerald^40^ was fused to the *C* terminus of the HERD. Finally, an *N*-terminal His tag followed by a TEV-cleavage site were added for purification. We named the final construct His–HERD-0–mGFP, or HERD-0–GFP for short. The constructs below are similar but with the HERD varied (Supplementary Table 1).

Expression of HERD-0–GFP in *E. coli* resulted in fluorescent intracellular foci (Fig. 2a,b; Supplementary Fig. 2), whereas, expression of mEmerald alone gave uniformly distributed fluorescence, indicating that protein condensation was specific to the HERD-0–GFP construct (Supplementary Fig. 3). However, western blotting showed that the majority of the *de novo* polypeptide was irreversibly aggregated (Supplementary Fig. 4), which we attributed to the strong (≤nM affinity) interactions between the HRs.^26,35^ Therefore, to weaken these CC interactions and the net PPIs, initially, we shortened the HRs from the standard 28 residues to 21 residues (Fig. 1e). By itself, this reduction in HR length produced polypeptides that did not differ significantly from the originally designed HERD-0–GFP in condensation or solubility (Supplementary Figs. 5 and 6). Therefore, we applied a combination of the following strategies: (1) further shortening the HRs; (2) mutating interfacial hydrophobic residues to alanine (Ala) (Fig. 1f); and (3) introducing overall helix-destabilising mutations^41^ outside of the hydrophobic interface (Fig. 1g). This gave the redesigns HERD-2.1– GFP through 2.4. The resulting constructs improved solubility while retaining protein condensation in cells (Fig 2a,b and Supplementary Figs. 7 and 8). However, destabilisation of the HRs beyond the recognised limit of CC formation—for instance, by reducing their lengths to two heptads or less, and completely disrupting core packing (HERD-2.5–GFP through 2.8, and HERD-Ctrl1–GFP and 2)^42^—resulted in the loss of protein condensation and gave largely dispersed and soluble constructs (Supplementary Fig. 9 – 12). At this stage different linkers were tested (HERD-3.1– GFP through 3.4), but this had no discernible impact on protein condensation (Supplementary Fig. 12).

**Figure 2:**
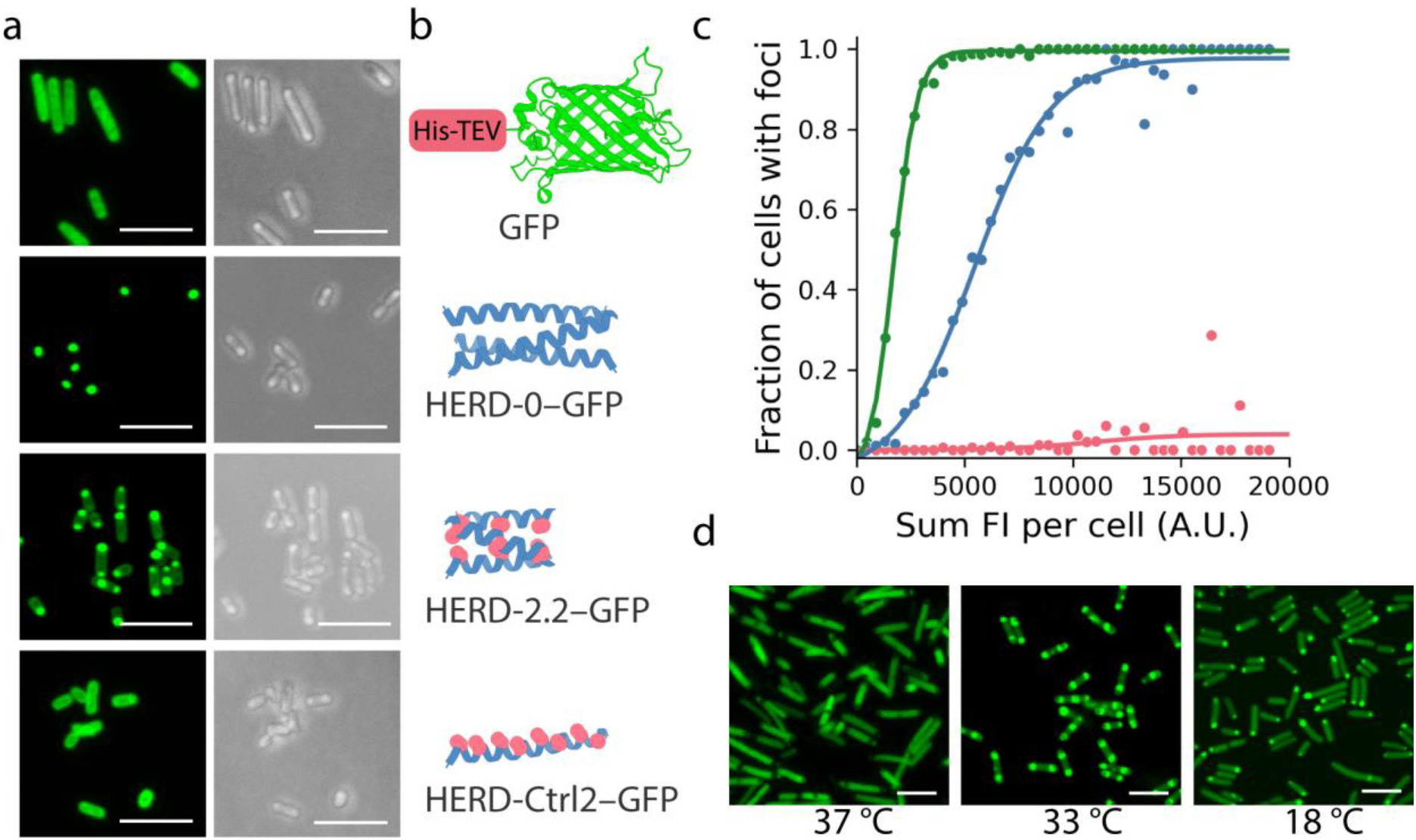
Weakening the designed helix-helix interactions leads to soluble protein that can condense in cells. **a & b,** Progression of the HERD designs visualised by light microscopy. **a,** Fixed-cell confocal microscopy images of *E. coli* cells expressing soluble GFP (His-TEV-GFP; control), the initial HERD-0–GFP design, the final variant HERD-2.2–GFP, and a control construct with a monomeric helical region HERD-Ctrl2–GFP (control). GFP fluorescence images (488 nm) are shown left (green) and brightfield transmission images on the right (grey). Scale bars, 5 μm. **b,** Cartoons of the helical regions of the HERD design, with Ala substitutions highlighted red. The linkers and GFP are mostly omitted for clarity. **c,** Automated image analysis of protein condensation in *E. coli* cells expressing HERD-0–GFP (green; n = 5782), HERD-2.2–GFP (blue; n = 5993), and HERD-Ctrl2–GFP (pink, n = 7923). *E. coli* cells were binned according to their total intra-cellular fluorescence (x-axis) and the fraction of cells identified as displaying intracellular foci (y-axis). *E. coli* cells were grown for 6 hours after induction at 18 °C and collected hourly for imaging and automated foci detection. **d,** Live-cell confocal microscopy images of HERD-2.2–GFP in *E. coli* at different growth temperatures. At 37 °C and 33 °C the formation of non-fluorescent inclusion bodies are visible by non-fluorescent cellular foci. Scale bars, 5 μm.

Automated image detection of foci in *E. coli* was used to quantify changes in cellular protein concentration and condensation (Fig. 2c and Supplementary Fig. 13). For HERD-0–GFP, this revealed condensates formed at low protein concentrations, suggesting aggregation. By contrast, HERD-2.2–GFP only formed condensates when a critical intracellular concentration was reached, indicative of threshold phase separation. Interestingly, this behaviour was also temperature dependent: in *E. coli* grown at 37 °C there was no observable condensation, while at lower temperatures (33 °C and 18 °C) enriched protein condensates were observed (Fig. 2d). Therefore, HERD-2.2–GFP was selected for further analysis. This has 2.5-heptad (17-residue) HRs with Ala residues at the ***a*** positions and isoleucine (Ile) at the ***d*** sites.

### Purified HERD-2.2–GFP phase separates *in vitro*

We purified the intact HERD-2.2–GFP protein for *in vitro* studies (Supplementary Fig. 14). Initially, different buffers, ionic strengths, and molecular crowders^43^ (*i.e.*, PEG 3350) were screened by brightfield microscopy to identify conditions for phase separation (Supplementary Fig. 15). We observed both general protein aggregation and potential liquid-liquid de-mixing, characterised by the formation of spherical macroscopic droplets (Fig. 3a and Supplementary Fig. 15). The optimal conditions for droplet formation were 4% PEG 3350 and 150 mM NaCl in Tris-HCl buffer (pH 7.5). Observations under these conditions by confocal microscopy revealed that the droplets were spherical and coalesced, indicative of liquid-like behaviour (Fig. 3b,c). Variable-temperature measurements showed that droplet formation occurred as the temperature was reduced from 40 °C to 5 °C (Supplementary Fig. 16, and Supplementary Video 1), and was reversible upon reheating (Supplementary Video 2). All of these properties are consistent with the formation of liquid condensates formed by LLPS. Also, we tested for any contribution of the *N*-terminal His-TEV tag: following TEV cleavage, the shortened HERD-2.2–GFP still underwent phase separation similar to the full-length protein, though it required slightly more molecular crowding agent, 10% PEG 3350, (Supplementary Figs. 17 and 18).

**Figure 3:**
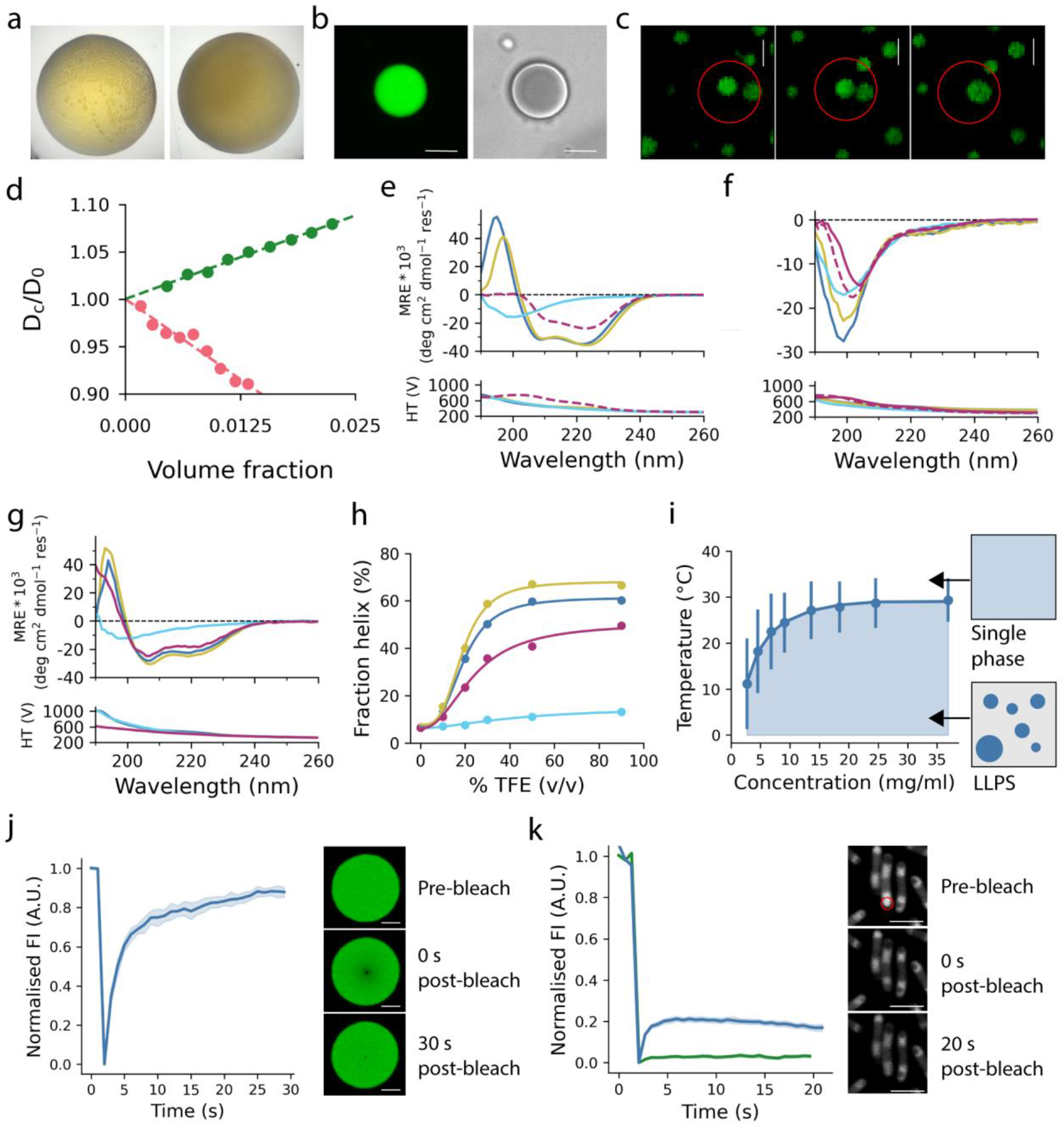
HERD-2.2–GFP forms de-mixed liquid droplets *in vitro* and in cells. **a,** Images of HERD-2.2–GFP showing macroscopic liquid de-mixing (left) and amorphous aggregation (right). **b,** Confocal microscopy of 1 mM HERD-2.2–GFP de-mixed droplets in 4% PEG 3350; and **c,** showing the coalescence of 2 such droplets circled in red. Scale bar, 5 μm. **d,** The dependence of D_C_/D_0_ (D_C_, collective diffusion coefficient; D_0_, free-particle diffusion coefficient) on protein volume fraction for GFP (green) and HERD-2.2–GFP (pink) measured by dynamic light scattering at 20 °C. **e – h**, CD data for the chemically synthesised HR and linker peptides. Key: HR1, blue; HR2, yellow; linker, teal; mixture, purple dashes; HR1–linker–HR2, purple solid. **e & f,** CD spectra for the HERD-0 (**e**) and HERD-2.2 (**f**) peptides at 500 μM (per peptide). **g,** CD spectra of HERD-2.2 peptides at 100 μM in 50% TFE, 5 °C. **h,** Fraction helix of HERD-2.2 peptides through a TFE titration, at 100 μM peptide, 5 °C. **i,** Phase diagram of 2 mM HERD-2.2–GFP from turbidity measurements in 4% PEG 3350. Bars indicate the difference between T_cloud_ and T_clear_. **j,** FRAP of HERD-2.2–GFP droplets *in vitro*. t_1/2_ = 1.54 ± 0.21 s. Shaded area represents the standard error (n = 13). Representative images of pre-bleach, post-bleach frame 1, and the final post-bleach frame shown alongside. Scale bar, 5 μm. **k,** FRAP of HERD-2.2–GFP (blue; n = 13) and HERD-0–GFP (green; n = 19) condensates in cells. t_1/2_ = 0.46 ± 0.11 s. Representative images of pre-bleach, post-bleach frame 1, and the final post-bleach frame shown alongside. Scale bar, 5 μm. Common conditions for all experiments: 125 mM NaCl, 20 mM Tris-HCl, pH 7.5.

To examine the self-interactions of the designed polypeptide further, we compared the diffusion (net) interaction parameter, *k*_D_, for GFP and HERD-2.2–GFP (Fig. 3d).^44^ Positive values of *k*_*D*_ indicate a protein with repulsive net interactions, while negative values indicate attractive PPIs. While GFP was slightly repulsive for itself (*k*_D_ = 3.5 ± 0.2), the fusion protein was overall attractive (*k*_D_ = -6.9 ± 0.6) consistent with its behaviour in cells and *in vitro*. Moreover, this *k*_D_ value is similar to those measured for other proteins that undergo LLPS.^45^ Therefore the designed polypeptide tag introduced attractive interactions to the slightly repulsive GFP molecule, making the fusion construct overall attractive for itself.

### Nascent helicity helps drive the condensation of HERD-2.2–GFP

To probe changes in secondary structure content of HERD-2.2–GFP, we followed de-mixing by circular dichroism (CD) spectroscopy. CD spectra were dominated by the β structure of GFP, and showed no detectible changes when recorded at different protein concentrations and temperatures (Supplementary Fig. 19). To investigate this further, we made the variants of the HR1 and HR2 sequences from our design trajectory and the linker sequence by solid-phase peptide synthesis (Supplementary Table 2; Supplementary Figs. 20 and 21). As expected, the CD spectra of the original HRs from the HERD-0 domain were highly α-helical, with intense minima at 208 nm and 222 nm (Fig. 3e; Supplementary Fig. 22); while the spectrum of the linker had a single minimum at 200 nm and low signal at 222 nm, indicating disorder. By contrast, for the HR variants of HERD-2.2—*i.e.*, with Ala at ***a*** positions, and truncated to 2.5 heptads— this helicity was lost, although some residual structure over the disordered linker peptide may be present (Fig. 3f; Supplementary Fig. 22). Mixing this HR1, HR2, plus linker combination did not induce further structure (Fig. 3f). Moreover, a chemically synthesised HR1–linker–HR2 peptide for HERD-2.2 appeared largely unstructured too, even in the presence of PEG (Supplementary Figs. 22 and 23).

These *in vitro* CD data were unexpected given our design hypothesis that α-helical domains drive PPIs and phase separation. However, it is possible that destabilised HRs still form nascent helices that associate weakly in the crowded environment of phase-separated droplets in cells.^46^ To investigate the potential for nascent helicity in the HERD-2.2 design, we recorded CD spectra in the presence of trifluoroethanol (TFE).^47^ In TFE titrations, the HR1 and HR2 peptides shifted to α-helical conformations, whereas the linker remained unstructured (Fig. 3g,h and Supplementary Fig. 24). Consistent with this, the titration for the synthetic HR1–linker– HR2 peptide led to less helix than observed for either of the HRs. These experiments indicate that, whilst largely unstructured in aqueous solution, the HRs of HERD-2.2 have the propensity for form α helices when the conditions are perturbed.

Encouraged by these data, we tested for nascent helicity and helix-helix interactions in cells by mutating HR1 and HR2 in the successful HERD-2.2–GFP background to knock out any such structure and interactions (controls 1 – 7, Supplementary Table 1). For instance, we replaced the remaining large hydrophobic Ile residues with Ala in a 3 heptad HERD background, to resemble a known monomeric α helix (HERD-Ctrl1–GFP & Ctrl2).^48^ To eliminate the amphipathicity of the HRs, we scrambled their sequences (HERD-Ctrl3–GFP). And, we introduced helix-breaking mutations into the HRs, *e.g.* proline (Pro, P) or glycine (Gly, G) at various positions of the heptad repeats (HERD-Ctrl4–GFP through Ctrl7). All of these redesigns showed significantly reduced or near-complete abolishment of protein condensation in cells (Fig. 2c and Supplementary Figs. 13 and 25). From these experiments, we posit that interactions between partially or transiently helical regions in the parent construct, HERD-2.2–GFP, are the most likely drivers of condensation.

### HERD-2.2–GFP undergoes LLPS *in vitro* and in cells

Next, we mapped the binodal phase boundary of expressed and purified HERD-2.2– GFP by measuring the cloud-point as a function of temperature and protein concentration (Supplementary Fig. 26). These experiments started with a single phase at higher temperature and measured changes in turbidity as the phases separated upon cooling. This returned an upper critical solution temperature (UCST) for an enthalpically driven phase transition (Fig. 3i). The process was reversible on heating with hysteresis between T_cloud_ and T_clear_ characteristic of protein LLPS. Further, the turbidity change accelerated with increased protein concentration consistent with faster nucleation.

To confirm the liquid nature of the condensates, we probed molecular diffusion within the droplets by fluorescence recovery after photobleaching (FRAP). First, droplets of purified HERD-2.2–GFP showed FRAP with a t_1/2_ of 1.54 s and near-complete recovery of signal, indicating highly mobile molecules (Fig. 3j). Next, we performed FRAP on the HERD-2.2–GFP condensates directly in *E. coli* cells. Here, we observed recovery of fluorescence with a similar rate to that measured *in vitro* (< 1 s; Fig. 3k). However, compared with the bulk *in vitro* experiments the amplitude of the final signal was considerably reduced (Fig. 3j,k). We attribute this to the confined system of the cell and, thus, the finite amount of fluorescent protein available to diffuse into the bleached region, which is large relative to the volume of the cell. This contrasts with the *in vitro* experiments where there is effectively an infinite amount of unbleached material to diffuse back into the bleached area. Nonetheless, in cells, asymmetrically bleached droplets nearly entirely re-equilibrated their fluorescence only 20 s after bleaching (Supplementary Fig. 27). Similar experiments with HERD-0–GFP condensates in cells showed no fluorescence recovery, and asymmetrically bleached droplets did not re-equilibrate their fluorescence between the bleached and non-bleached areas (Fig. 3k and Supplementary Fig. 26). Thus, our design process progressed from insoluble CC-based constructs (with HERD-0) to biomolecular condensates with dynamic, liquid properties (with HERD-2.2) both *in vitro* and in cells.

### HERD-2.2 condensates can be functionalised in cells

Finally, we tested the HERD-2.2 polypeptide as a component for designing functional membraneless organelles with alternate client proteins. Initially, we swapped mEmerald for mCherry to give two fluorescent constructs HERD-2.2–GFP and HERD-2.2–mCherry (Fig. 4a). When co-expressed in *E. coli*, these co-localised into the same condensates (Fig. 4a and Supplementary Fig. 28). Next, we replaced the fluorescent proteins with the enzymes tryptophanase (TnaA) and flavin-containing monooxygenase (FMO) to give HERD-2.2–TnaA and HERD-2.2–FMO (Supplementary Fig. 29). TnaA and FMO catalyse the two-step conversion of tryptophan to indigo (Fig. 4b), which we sought to test in the HERD-based system.^49^ Purified, His-tagged HERD-2.2–TnaA formed de-mixed liquid droplets similar to HERD-2.2–GFP, while HERD-2.2–FMO did not undergo phase separation *in vitro* (Fig. 4c and Supplementary Fig. 30). We attribute this to TnaA and GFP having very similar calculated net charges (both -6 at pH 7.5), whereas FMO is highly negatively charged (net charge -21 at pH 7.5). Again, this indicates that it is the net PPIs made by whole construct and not just by the *de novo* polypeptide that leads to condensation.

**Figure 4:**
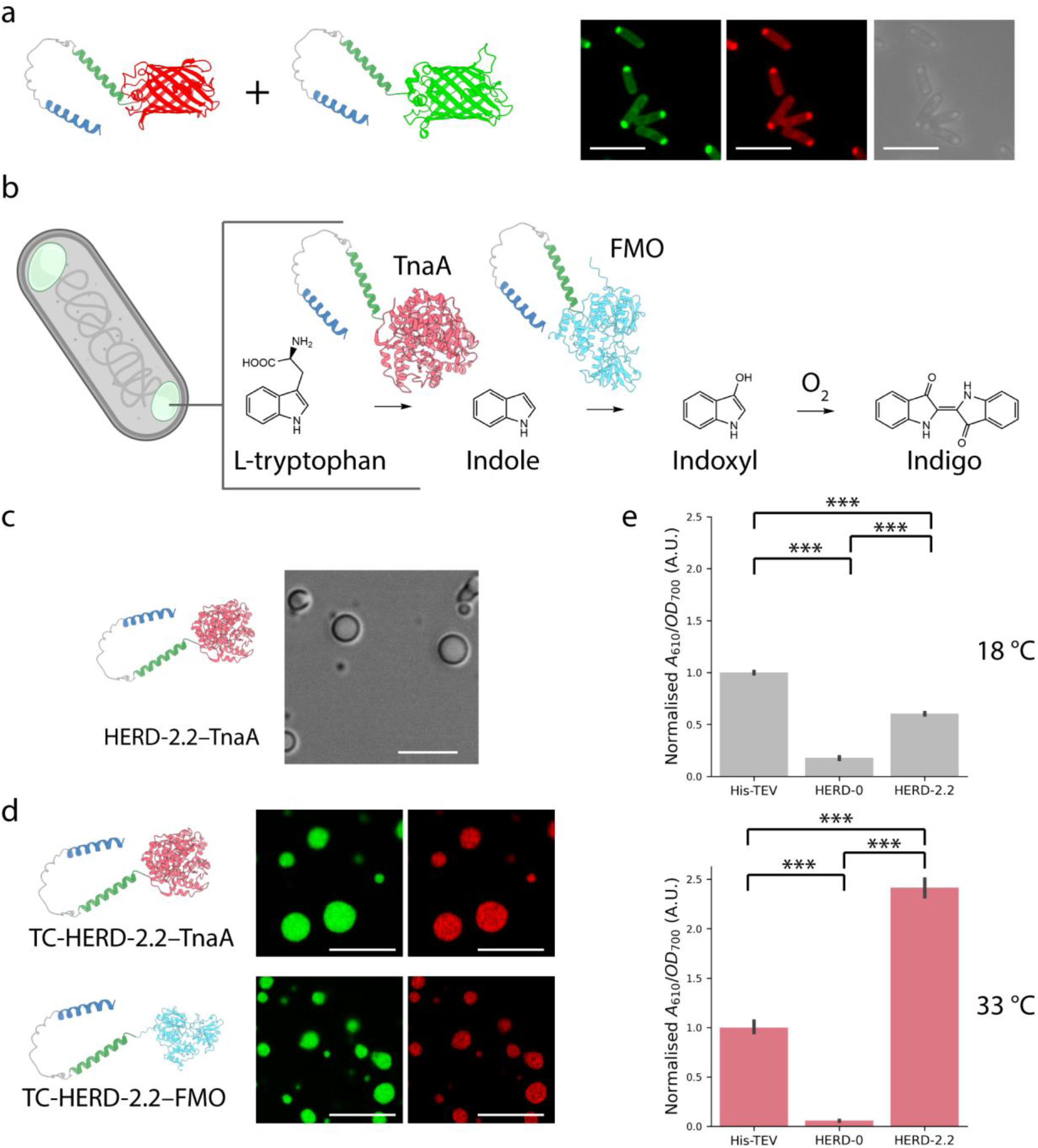
HERD-2.2-tagged enzymes form functional membraneless organelles. **a,** Confocal microscopy images of HERD-2.2–GFP and HERD-2.2–mCherry co-expressed in *E. coli.* GFP fluorescence at 488 nm (green), mCherry fluorescence at 561 nm (red), and brightfield transmission (grey). Scale bar, 5 μm. **b,** Schematic for the in cell colocalization of TnaA and FMO using the HERD-2.2 polypeptide and the subsequent enzymatic production of indigo dye. **c,** Brightfield image of HERD-2.2–TnaA droplets *in vitro*. Scale bar, 5 μm. Conditions: 200 mM NaCl, 8% PEG 3350, 40 mM Bis-Tris pH 6, 120 μM HERD-2.2–TnaA. **d,** Confocal microscopy images of de-mixed droplets *in vitro* formed by mixing HERD-2.2–GFP plus TC-HERD-2.2–TnaA (upper), and HERD-2.2–GFP plus TC-HERD-2.2–FMO (lower), with fluorescent reporters GFP (green) and TC-ReAsH at 561 nm (red). Scale bar, 5 μm. Conditions: 125 mM NaCl, 4% PEG 3350, 20 mM Tris pH 7.5, 2 mM HERD-2.2–GFP, 100 μM HERD-2.2–TnaA or HERD-2.2–FMO. **e,** Normalised indigo production in cells co-expressing fusions of TnaA and FMO plus the GFP scaffold at 18 °C (grey) and 33 °C (pink). Error bars represent the standard error (n = 3 for all conditions). P value < 0.001 for all groups measured by one way ANOVA and pairwise TukeyHSD *post hoc* test.

As HERD-2.2–FMO did not undergo LLPS in isolation, we tested if HERD-2.2–GFP could facilitate the co-condensation of the FMO and TnaA constructs. First, we confirmed that HERD-2.2–GFP droplets *in vitro* could recruit both HERD-2.2–FMO and HERD-2.2–TnaA separately by adding tetra-cysteine motifs into the flexible linkers (giving TC–HERD-2.2–FMO and TC–HERD-2.2–TnaA, Supplementary Table 1) and subsequently labelling with TC-ReAsH II^50^ (Fig. 4d and Supplementary Figs. 31 and 32). Assuming that both tagged proteins would co-condense with HERD-2.2 droplets, we tested the effect of enzyme colocalization directly in cells. HERD-2.2–TnaA and HERD-2.2–FMO fusions were co-expressed under the arabinose promotor on a low copy number vector to control their expression, while HERD-2.2–GFP was expressed from the T7 promotor to generate the de-mixed compartments. Initially, indigo production in *E. coli* was measured for cells grown at 18 °C (Fig. 4e, Supplementary Fig. 33). Under these conditions the system performed worse than free enzymes expressed with only *N*-terminal His-TEV tags. This stands to reason as FRAP performed in cells at 18 °C indicated that these condensates were less liquid-like and possibly amorphous aggregates (Supplementary Fig. 34). By contrast, at 33 °C—*i.e.*, closer to where HERD-2.2–GFP condensates were identified as forming a dense liquid phase (Fig. 3k and Supplementary Fig. 27)—the combination of the three constructs produced nearly 2.5 times as much indigo as the free enzymes, suggesting that efficiency of the enzyme cascade was improved by co-condensation to a liquid-state. This increase in indigo production was still significant after normalisation for enzyme expression levels (Supplementary Fig. 35). As a control, under both growth conditions, the HERD-0–GFP showed significantly reduced indigo production compared the free enzymes. This suggests that the HERD-0-based design simply sequesters the enzymes and makes them inaccessible to substrate.

## Conclusion

In summary, we have designed a polypeptide tag that can be fused to client proteins enabling the resulting fusions to undergo phase separation *in vitro* and in living cells. Rather than designing geometrically defined and rigid proteins, we focused on creating constructs that are reminiscent of IDRs and promote long range disorder to create macroscopic condensates with desired physical properties. The polypeptide is designed from first principles by concatenating two different helical oligomerisation domains *via* an intrinsically disordered sequence to give ≈100-residue sequence. When appended as an *N*-terminal tag to a green fluorescent protein this produces insoluble aggregates in cells. However, the assemblies can be rendered soluble by destabilising the helical regions and weakening their interactions. Notably, only a handful of designs had to be screened to identify sequences with the desired characteristics, and in the final design, condensate formation depends on protein concentration and temperature indicative of reversible liquid-liquid phase separation (LLPS). Through a series of *in vitro* and in-cell experiments using soft-matter physics and biophysical methods, we show that the protein condensates are highly dynamic and behave like de-mixed liquid droplets. Finally, we demonstrate that the client fluorescent protein can be substituted by an enzyme and that the resulting droplets can recruit another tagged enzyme that by itself does not undergo LLPS. In cells, these dual-enzyme organelles give increased activities—albeit modest—over the expressed soluble enzymes to highlight the potential utility of this system.

Others have reported the successful engineering of natural condensing proteins in cells to explore the potential of MLOs in synthetic biology.^12,15,17,18,51,52^ Recent examples have demonstrated the capability for condensates to augment cells with artificial functions approaching the complexity of natural organelles,^14,53^ and to generate compartments that can be modulated through rational changes to their scaffold proteins.^16^ Engineered condensates have also incorporated designed enzymatic reactions, creating functional MLOs.^54^ In addition to colocalizing client proteins, our system has a potentially useful thermo-switchable behaviour that allows induction and dissolution of protein condensates *in vitro* and in cells by modifying the conditions or the cell-growth temperature. Furthermore, the material properties of these condensates can be varied with temperature, switching from gel-like condensates to liquid-like droplets. This directly affects the efficiency of the co-condensed enzyme cascade, as measured in our system by the production of indigo. This temperature-dependent switching of phase behaviour potentially permits the control of protein condensation and function using a simple control mechanism.

Overall, we anticipate that our *de novo* designed polypeptide tag will provide a valuable tool for studying biomolecular condensation, and for developing membraneless organelles in synthetic or natural biological systems both *in vitro* and within cells. Furthermore, the relative simplicity of our designs and the ease with which they can be redesigned to access soluble, dynamic condensed, and aggregated states should allow them to be adapted readily for other experiments and applications.

## Acknowledgements

ATH and DNW are funded by the University of Bristol through the Max Planck-Bristol Centre for Minimal Biology. AR is funded by the Leverhulme Trust through a grant to JJM and DNW (RGP-2021-049). RO was funded through a European Union’s Horizon 2020 research and innovation programme Marie Skłodowska-Curie grant (NovoFold No. 795867). We thank the University of Bristol School of Chemistry Mass Spectrometry Facility for access to the EPSRC-financed Bruker Ultraflex MALDI-TOF/TOF instrument (EP/K03927X/1), BrisSynBio for access to peptide synthesizers (BB/L01386X/1), and the Wolfson Bioimaging Facility for their assistance in this work. We thank Prof Chong Zhang of Tsinghua University for providing a BL21 (DE3) ΔtnaA *E. coli* strain. We thank Dr Matt Lee for the pDIC bacterial vectors. We thank Dr Peter Wilson (BioSuite, University of Bristol) for use of the Formulatrix crystallisation hotel, and Dr Alessandro Strofaldi for assistance with the soft-matter experiments.

## Materials & Methods

### Materials

All chemicals and biological materials were obtained from commercial suppliers. Escherichia coli BL21(DE3), Q5 DNA polymerase, T4 DNA ligase, and restriction enzymes were purchased from New England BioLabs. Genes were ordered as g-blocks from IDT, primers were ordered from Eurofins Genomics. Anti-His primary antibody (H1029) was purchased from Sigma Aldrich, HRP conjugated secondary antibody (J1430) was purchased from Invitrogen. LB broth (Lennox) was purchased from Sigma Aldrich.

### Calculation of protein pI and net charge

Protein physical and chemical parameters were calculated from the primary amino acid sequence using the ExPASy ProtParam tool.^1^

### Protein expression

The genes encoding the HERDs and enzymes (FMO, TnaA) were codon optimized for expression in *E. coli* and ordered from IDT as g-blocks. Constructs were cloned into pET38a(+) derivative vectors (pDICa - ampicillin selection marker or pDICc - chloramphenicol selection marker; Supplementary Fig. 36) kindly donated by M. Lee, using XbaI and NdeI restriction sites and T4 DNA ligase.

Plasmids (25 ng) were transformed into *E. coli* BL21(DE3) competent cells by heat shock and plated on LB agar plates supplemented with appropriate antibiotics (100 μg/ml ampicillin, 25 μg/ml chloramphenicol). Following overnight incubation at 37 °C, a single colony was used to inoculate 5ml LB and grown overnight (37 °C, 200 rpm). Fresh LB was inoculated 1:100 from the overnight culture and grown to OD_600_ = 0.4 - 0.6 (37 °C, 200 rpm). Protein expression was then induced using 400 μM IPTG (vectors with the T7 promotor) or 0.2% D-arabinose (vectors with the arabinose promotor).

### In-cell confocal microscopy

For confocal microscopy, 50 ml of LB was inoculated from the overnight culture. After induction of protein expression, cultures were grown at 18 °C, shaking at 200 rpm typically for 5 hours. 1 ml of culture was collected and cells pelleted by centrifugation (3000 xg, 3 minutes). For fixed cells, pellets were washed 3 times in PBS, before fixing by incubating in 1 ml of 2% paraformaldehyde in PBS for 15 minutes at room temperature. Pellets were washed a further 3 times in PBS before resuspending in 50 μl PBS. Fixed cells were mounted in ProLong Diamond Antifade Mountant (Invitrogen). For live cell imaging, cells were grown as described above, with variable growth temperatures (18 – 37 °C) after induction. 1 ml of culture was collected by centrifugation and immediately resuspended in 50 μl PBS. 15 μl of cell suspension was sealed onto a glass slide under a coverslip with nail polish to prevent evaporation and imaged immediately. Confocal images were collected using a Leica SP5II microscope using a 63x objective lens, running Leica LAS X. Fixed cell images are represented as maximum intensity projections, assembled in ImageJ.

### Western blotting

For western blotting, 50 ml of LB was inoculated from the overnight culture and grown at 18 °C, 200 rpm. Pellets were collected by centrifugation (3000 xg, 10 mins) after normalisation to cell density (OD_600_). Pellets were lysed by resuspension in BugBuster lysis buffer (Millipore) with Benzonase nuclease (Millipore) and incubated at 37 °C for 30 minutes. Suspensions were then snap frozen 3 times in liquid nitrogen to ensure complete cell lysis. For separation of cellular soluble and insoluble fractions, suspensions were centrifuged (18000 xg, 20 minutes). The supernatant (soluble fraction) was removed and the pellet (insoluble fraction) resuspended in an equal volume of BugBuster. For SDS-PAGE, samples were mixed with appropriate volumes of reducing SDS loading dye and boiled at 95 °C for 5 – 10 minutes. 6 μl of sample was loaded alongside 6 μl of colour pre-stained protein standard, broad range (NEB) onto 12% acrylamide/bis-acrylamide (29:1) gels and run at 180 V for 1 hour, or until the loading dye reached the bottom of the gel. For western blotting, proteins were transferred onto a 0.2 μM PVDF membrane (Cytiva) using Power Blotter 1-Step™ Transfer Buffer (Invitrogen) for 10 minutes at 1.3 A. Membranes were blocked in 4% skimmed milk powder with 0.1% Tween-20 in PBS for 30 minutes with gentle rocking. Membranes were then incubated with anti-His primary antibody 1:5000 in 4% milk in PBS-T (Sigma) for 2 hours. Membranes were washed 3 times for 5 minutes in PBS-T, before adding the HRP conjugated secondary antibody 1:10000 in 4% milk in PBS-T (Invitrogen) for 1 hour. Membranes were washed a further 3 times for 5 minutes in PBS-T, before adding 2 ml of SuperSignal™ West Pico Plus chemiluminescent substrate (Thermo), and incubating for 1 minute before imaging using a G:Box Chemi-XT4 chemiluminescent imager (SynGene) for the desired interval.

### Automated image analysis

Brightfield and fluorescent microscopy images of *E. coli* were quantified using the ModularImageAnalysis (MIA; v0.21.11) plugin for Fiji.^2-5^ Prior to detection of *E. coli*, brightfield images stacks were normalised using sliding paraboloid background subtraction (radius = 10 px). From these, single slices chosen for optimal feature contrast were extracted using a modified version of the Stack Focuser ImageJ plugin.^6^ The focused images were then intensity normalised and subject to further background correction by pixelwise division with 2D Gaussian-filtered (sigma = 10 px) variants of the same images. The corrected brightfield images were down-sampled 2x in XY before being passed to the StarDist Fiji plugin for detection of *E. coli*,^7-9^ using a model trained on the DeepBacs *E.coli* dataset.^10^ To account for overlap between adjacent cells detected via StarDist, binary images showing detected cells were created and resegmented using the distance-based watershed transform. Final *E. coli* detections were obtained from the segmented images using connected components labelling. Foci were detected in maximum intensity z-axis projections of fluorescent image stacks. These images were passed through a 2D top-hat filter (radius = 5 px) to remove general cell background intensity. The images were then converted to binary maps using a fixed global intensity threshold and adjacent foci separated using another distance-based watershed transform. Markers for the watershed transform were acquired using TrackMate’s LoG spot detector.^11^ This detector convolves the image with a Laplacian of Gaussian kernel to enhance spot-like features of a specific size (radius = 4 px) and detects foci as features in the convolved image brighter than a set threshold. Foci were detected from the segmented images using connected components labelling.^12^ Number, area and fluorescent intensity statistics for each measured cell and focus were measured and exported as a single Excel spreadsheet for downstream analysis.

### Protein purification

For protein purification, 1 to 12 l of LB was inoculated 1:100 from an overnight culture and grown at 18 °C, shaking at 200 rpm. Cell pellets were resuspended in buffer containing 500 mM NaCl, 20 mM Tris-HCl pH 7.5, 2 M urea, 50 mM imidazole, 1 tablet cOmplete protease inhibitor (Roche), and lysed by sonication on ice (3 s on, 1 s off, 70% amplitude, 15 minutes). The lysate was centrifuged (18000 xg, 20 minutes) and the supernatant filtered through a 0.2 μM filter to clarify. Protein purification was performed using an Äkta Pure (Cytiva) at 4 °C, with chromatograms monitored at 280 nm. The clarified lysate was applied to a HisTrap HP (Cytiva) immobilised metal affinity chromatography (IMAC) column, pre-equilibrated in 500 mM NaCl, 20 mM Tris-HCl pH 7.5, 2 M urea, 50 mM imidazole. The column was washed until A_280_ was re-stabilised (typically 3 – 4x the column volume), before eluting the bound protein with a gradient of imidazole (50 – 500 mM). Recombinant protein was further purified by size exclusion chromatography using a HiLoad 16/600 Superdex 200 pg exclusion column (Cytiva) with a flow rate of 1 ml/min. Size exclusion was performed using a 20 mM Tris-HCl pH 7.5, 2 M urea running buffer and elution monitored by A_280_. Protein fractions were identified by SDS-PAGE and the relevant fractions pooled. Protein samples were finally desalted using a HiPrep 26/10 desalting column (Cytiva) into 20 mM Tris-HCl pH 7.5, aliquoted, flash frozen and stored at -70 °C.

### Circular-dichroism spectroscopy

Circular-dichroism (CD) data were collected on a JASCO J-810 or J-815 spectropolarimeter fitted with a Peltier temperature controller (Jasco UK). Full spectra were measured between 190 and 260 nm with a 1 nm step size, 100 nm·min^-1^ scanning speed, 1 nm bandwidth and 1 second response time. Spectra were measured at 5 °C unless otherwise stated. Baselines recorded using the same buffer, cuvette and parameters were subtracted from each dataset. For experiments in TFE, the protein in buffer was mixed with neat TFE to produce the stated concentrations. The spectra were converted from ellipticities (deg) to mean residue ellipticities (MRE, (deg.cm^2^.dmol^-1^.res^-1^)) by normalizing for concentration of peptide bonds and the cell path length using the equation:

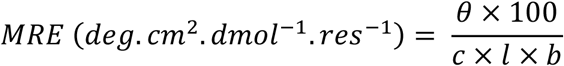

Where the variable *θ* is the measured difference in absorbed circularly polarized light in millidegrees, *c* is the millimolar concentration of the specimen, *l* is the path-length of the cuvette in cm and *b* is the number of amide bonds in the polypeptide, for which the *N*-terminal acetyl bond was included but not the *C*-terminal amide. Peptide concentration was determined at 280 nm (ε_280_(Trp) = 5690 cm^-1^, ε_280_(Tyr) = 1280 cm^-1^)^13^ (for the peptides 1 – 9) or by measuring the peptide bond at 214 nm^14^ (for the peptide 10) using a Nanodrop 2000 (Thermo) spectrometer. Fraction helix (%) was calculated from MRE at 222 nm using the following equation:

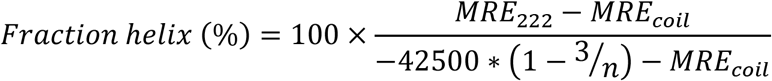

Where *MRE*_*coil*_ = 640-45*T*; *T* is the temperature in °C; and *n* is the number of amide bonds in the sample (including the C-terminal amide).^15^

### Peptide synthesis

Solid-phase peptide synthesis (SPPS) reagents were purchased from Cambridge Reagents with the exception of N,N’-diisopropylcarbodiimide (DIC) purchased from Carbosynth. Rink amide MBHA resin and Fmoc-protected amino were purchased from Merck. SPPS was performed on a Liberty Blue automated peptide synthesizer (CEM) with inline UV monitoring. All peptides were synthesized as the *C*-terminal amide on Rink amide MBHA resin, with DIC/Oxyma as the coupling reagents. Fmoc was removed using 20% v/v morpholine:dimethylformamide (DMF). All peptides were *N*-terminally acetylated through treatment with pyridine (0.5 mL) and acetic anhydride (0.3 mL) in DMF (9.2 mL) and shaking at room temperature (rt) for 20-60 minutes. Peptides were cleaved from the resin with addition of 95:2.5:2.5 v/v trifluoroacetic acid (TFA):H2O:triisopropylsilane and shaking at room temperature for 3 hours. Following collection of the cleavage solution, TFA was evaporated under a N_2_ stream followed by precipitation with ice cold diethyl ether. Precipitates were collected by centrifugation and dissolved in 50:50 v/v acetonitrile (MeCN):H2O. Crude peptides were lyophilized to yield a white or off-white powder

### Peptide purification

Peptides were purified by reverse phase HPLC on a Phenomenex Luna C18 stationary phase column (150 × 10 mm, 5 μM particle size, 100 Å pore size) using a preparative JASCO HPLC system. Crude peptide was dissolved at 3 – 5 mg/mL in 0 – 20% v/v acetonitrile with 0.1% TFA. A (0 – 20) – 100% gradient of acetonitrile with 0.1% TFA over 30 – 45 minutes was used to separate the target peptide. Chromatograms were monitored at wavelengths of 220 and 280 nm. The identities of the peptides were confirmed using mass spectrometry. Peptide purities were determined using a JASCO analytical HPLC system, fitted with a reverse-phase Kinetex® C18 analytical column (100 × 4.6 mm, 5 μm particle size, 100 Å pore size). Fractions containing pure peptide were pooled and lyophilised.

### Mass spectrometry

Matrix-assisted laser desorption/ionization–time of flight (MALDI-TOF) mass spectra were collected on a Bruker UltraFlex MALDI-TOF mass spectrometer operating in positive-ion reflector mode. Peptides were spotted on a ground steel target plate using α-cyano-4-hydroxycinnamic acid dissolved in 1:1 acetonitrile:H_2_O as the matrix. Masses quoted are for the monoisotopic mass as the singly protonated species. Full electrospray ionization (ESI) MS spectra were acquired on a Synapt G2S (Waters) mass spectrometer equipped with an IMS-Q-TOF analyser and using an Advion Nanomate for robot chip-based nanospray ionization in positive mode. 5 μl of a 50 μM peptide solution in 1:1 acetonitrile:H_2_O were generally injected for the analysis. Masses quoted are for the deconvoluted monoisotopic mass.

### TEV cleavage

Cleavage by TEV protease was performed using ProTEV Plus (Promega) with 1 mM DTT, 0.5 mM EDTA, 18 mg of HERD-2.2–GFP, and 200 units of ProTEV Plus in a 12 ml reaction volume. The reaction was incubated overnight at 30 °C. The cleaved protein was purified by application to a HisTrap HP column and collection of the flow-through. Cleavage was confirmed by SDS-PAGE and staining using Coomassie-blue.

### Dynamic light scattering

For DLS measurements, the proteins were purified as mentioned previously and desalted using a HiLoad 16/600 Superdex 200 pg column (Cytiva) with 20 mM Tris-HCl pH 7.5 buffer as an eluent the day before the experiment. Buffers were filtered through Anatop 0.02 μm filters (Whatman) were used for preparation of different protein concentrations. On the day of the experiment, the proteins were concentrated to 15 – 30 mg/ml concentration using Amicon Ultra Centrifugal filters (Merck) via short (2 – 5 min) cycles at the speed ≤ 3000 xg at 20 °C, and then centrifuged for 60 – 90 minutes at 17,000 ×g at room temperature to remove any pre-formed aggregates in solution.

An ALV/CGS-3 goniometer with a HeNe laser operating at a wavelength of 632.8 nm, an optical fibre based detector and an ALV/LSE-5004 Light Scattering Electronics and Multiple Tau Digital Correlator were used for DLS measurements. The temperature was kept constant at 20 °C during data acquisition using a Thermo Scientific DC30-K20 water bath connected to the instrument and measured with a Pt-100 probe immersed into the index matching fluid vat. DLS measurements were carried out for 30 – 60 minutes at a scattering angle of 90 °C at each protein concentration. The protein concentration was determined for the sample after the last measurement using Cary-100 (Agilent) UV-Vis spectrometer based on the extinction coefficients calculated by the ExPASy Server.^16^

Volume fraction is calculated using the expression *c* = Φ×*n* where *c* is the concentration in mg ml^-1^, Φ is the volume fraction and *n* is the partial specific volume equal to 7.266 10^−4^ and 7.326 10^−4^ ml·mg^-1^ for HERD-2.2–GFP and GFP, respectively, as calculated using sedfit software.^17^

### Cloud-point measurements

Measurement of the binodal phase boundary was performed in a Perkin Elmer Lambda 35 UV/Vis spectrophotometer with a temperature controlled cuvette holder regulated by an external circulating water bath. Measurements were performed at 125 mM NaCl, 4% PEG 3350, and 20 mM Tris-HCl pH 7.5, with varying concentrations of HERD-2.2– GFP (2.7 – 37 mg/ml). Samples were filtered using a 0.2 μM filter and incubated at 40 °C in an incubator to maintain a single phase prior to measurement. For each sample concentration, solution temperature was measured using a thermocouple in the reference cuvette. Phase separation was monitored by transmission (%T) at 600 nm as the temperature was decreased from 40 °C to 5°C and T_cloud_ identified as the 50% transmission point. After %T stabilised, the temperature was returned to 40 °C and T_clear_ identified as the 50% transmission point. The threshold temperature for LLPS at that concentration was calculated as the mean of T_cloud_ and T_clear_.

### Fluorescence recovery after photobleaching

Fluorescence recovery after photobleaching (FRAP) was performed using a Leica SP8 AOBS confocal with a 65 mW Ar laser exciting at 488 nm. For each bleaching measurement 3 images were taken before bleaching and the mean intensity recorded as the pre-bleach fluorescence intensity. Bleaching was performed using a 100 ms (*in vitro*) or 1 ms (in cell) laser burst at 40% laser power, followed by imaging every 0.65 s for 20 – 30 s to record fluorescence recovery. For each bleaching measurement, recovery was normalised relative to the mean fluorescence intensity before bleaching (normalised to 1), and the minimum fluorescence intensity measured immediately after bleaching (normalised to 0) to allow comparison between different bleaching experiments. For *in vitro* measurements, de-mixed droplets were placed on a clean glass slide and covered with a cover slip before imaging. *In vitro* conditions were 33 mg/ml HERD-2.2–GFP, 125 mM NaCl, 20 mM Tris-HCl pH 7.5, and 4% or 10% PEG 3350. For in-cell measurements, FRAP was performed on live *E. coli* cells prepared as described under *in-cell confocal microscopy*. Normalised FRAP data was fitted in OriginPro to an exponential model *f*(t)=A.(1-e^-τ.t^), where A is the plateau intensity, τ is the fitted parameter, t is the time after bleaching. Half-lives were determined using the formula: t_1/2_ = ln(0.5)/τ.

### TC-ReAsH II labelling

Tetra-cysteine (TC) tagged proteins (CCPGCC) were site-specifically labelled using the TC-ReAsH II TC detection dye (Invitrogen). TC-HERD-2.2–TnaA and TC-HERD-2.2–FMO (100 μM) were incubated separately with 1 μM TC-ReAsH II and 1 mM TCEP for 1 hour, prior to mixing with 1 mM HERD-2.2–GFP. The mixture was phase separated by addition of buffer containing 8% PEG 3350, 250 mM NaCl, and 20 mM Tris-HCl pH 7.5 and droplets formed imaged at 488 nm (GFP) and 561 nm (TC-ReAsH) on a Leica SP8 confocal microscope. Fluorescence intensity was compared between droplets containing TC tagged enzymes and those containing no TC tags to measure specific enrichment due to the TC tagged enzymes.

### In-cell indigo production

Indigo production in cells expressing HERD-tagged TnaA and FMO was performed using Δtnaa BL21 (DE3) *E. coli* generously provided by Dr Chong Zhang^18^. Δtnaa *E. coli* was co-transformed with a vector encoding the relevant HERD-GFP protein under the control of the T7 promoter (AmpR), and a second duet-style expression vector encoding both the relevant HERD-TnaA and HERD-FMO proteins under the control of the arabinose promoter (CmR). 50 ml of LB was inoculated 1:100 with overnight culture and grown at 37 °C to OD_700_ = 0.4 – 0.6. Cell density was measured using OD_700_ to avoid discrepancies due to the absorbance spectrum of indigo.^19^ Cultures were then induced with 400 μM IPTG and 0.2% D-arabinose and grown at 18 °C or 33 °C, shaking at 200 rpm. For indigo measurements, 2 ml of culture was collected 22 hours after induction and pelleted by centrifugation (3000 xg, 5 minutes). Indigo concentration was measured by resuspension of cell pellets in N-methyl-2-pyrrolidone (NMP) and sonication to dissolve the indigo. Solutions were centrifuged (13000 xg, 3 mins) to remove cell debris, and indigo concentration measured by absorbance at 610 nm on a Perkin Elmer Lambda 25 UV/Vis spectrophotometer. To compare expression of TnaA and FMO quantitatively, samples were taken for western blotting as described above. Western blot samples were collected at the same time as indigo samples (22 hours after induction). Samples for western blotting were normalised against OD_700_ to collect equal cell numbers and western blotted against the His epitope labelled enzymes. Quantification of enzyme expression from western blots was performed in Image Studio Lite against triplicate cultures after setting the background intensity manually to an area with no intensity, to avoid interference from the co-expressed GFP.

