## Supplementary Material for "Assembling membraneless organelles from *de novo* designed proteins"

| HERD # | Helical region 1 (HR1)<br>*gabcdef gabcdef gabcdef gabcdef* | Linker | Helical region 2 (HR2)<br>*cdefgab cdefgab cdefgab cdefgab* |
|---|---|---|---|
| 0 | G EIAAIKE EIAAIKE EIAAIKW EIAAIKE G | ASPEPQPKPSGDPQSKQTPEPSRSQ | G EIQEQLK EIQEQLK EIQWQLK EIQEQLK G |
| 1.1 | G EIAAIKE EIAAIKW EIAAIKE G | ASPEPQPKPSGDPQSKQTPEPSRSQ | G EIQEQLK EIQWQLK EIQEQLK G |
| 2.1 | G EAAAIKE EAAAIKW EAAAIKE G | ASPEPQPKPSGDPQSKQTPEPSRSQ | G EIQEQAK EIQWQAK EIQEQAK G |
| 2.2 | G IKE EAAAIKW EAAAIKE G | ASPEPQPKPSGDPQSKQTPEPSRSQ | G QAK EIQWQAK EIQEQAK G |
| 2.3 | G EISSIKE EISSIKW EISSIKE G | ASPEPQPKPSGDPQSKQTPEPSRSQ | G EIQEQLK EIQWQLK EIQEQLK G |
| 2.4 | G EISSIKE EISSIKW EASSIKE G | ASPEPQPKPSGDPQSKQTPEPSRSQ | G EIQEQLK EIQWQLK EIQEQAK G |
| 2.5 | G EAAAIKW EAAAIKE G | ASPEPQPKPSGDPQSKQTPEPSRSQ | G EIQWQAK EIQEQAK G |
| 2.6 | G EASSIKE EASSIKE EASSIKW EASSIKE G | ASPEPQPKPSGDPQSKQTPEPSRSQ | G EIQEQAK EIQEQAK EIQWQAK EIQEQAK G |
| 2.7 | G EISSIKE EASSIKW EASSIKE G | ASPEPQPKPSGDPQSKQTPEPSRSQ | G EIQEQLK EIQWQAK EIQEQAK G |
| 2.8 | G EASSIKE EASSIKW EASSIKE G | ASPEPQPKPSGDPQSKQTPEPSRSQ | G EIQEQAK EIQWQAK EIQEQAK G |
| 3.1 | G IKE EAAAIKW EAAAIKE G | ASPEPQPKPSGDPQSKQTPEASPRPQ<br>PEPSGKPQSEQTPKASPEPQPKPS | G QAK EIQWQAK EIQEQAK G |
| 3.2 | G IKE EAAAIKW EAAAIKE G | ASESQKPS | G QAK EIQWQAK EIQEQAK G |
| 3.3 | G IKE EAAAIKW EAAAIKE G | ASEPKQSDPKGDPRSEQKPEKSESR | G QAK EIQWQAK EIQEQAK G |
| 3.4 | G IKE EAAAIKW EAAAIKE G | ASPTSYPAPSWAPQYLQTPGPSVSQ | G QAK EIQWQAK EIQEQAK G |
| Ctrl1 | G EAAAAKE EAAAAKW EAAAAKE G | ASPEPQPKPSGDPQSKQTPEPSRSQ | G EIQEQAK EIQWQAK EIQEQAK G |
| Ctrl2 | G EAAAAKE EAAAAKW EAAAAKE G | ASPEPQPKPSGDPQSKQTPEPSRSQ | G EAQEQAK EAQWQAK EAQEQAK G |
| Ctrl3 | G KEI EAAAIKW EAAAKIE G | ASPEPQPKPSGDPQSKQTPEPSRSQ | G QAK IEQWQAK EEQIQAK G |
| Ctrl4 | G PKE EAAAPKW EAAAPKE G | ASPEPQPKPSGDPQSKQTPEPSRSQ | G QAK EPQWQAK EPQEQAK G |
| Ctrl5 | G GKE EAAAGKW EAAAGKE G | ASPEPQPKPSGDPQSKQTPEPSRSQ | G QAK EGQWQAK EGQEQAK G |
| Ctrl6 | G IKE EAPAIKW EAPAIKE G | ASPEPQPKPSGDPQSKQTPEPSRSQ | G QAK EIQPQAK EIQEPAK G |
| Ctrl7 | G IKE EAGAIKW EAGAIKE G | ASPEPQPKPSGDPQSKQTPEPSRSQ | G QAK EIQGQAK EIQEGAK G |
| TC-2.2 | G IKE EAAAIKW EAAAIKE G | ASPEPQPKPSCCPGCCQTPEPSRSQ | G QAK EIQWQAK EIQEQAK G |

**Supplementary Table S1: Sequences of HERD tags.** Sequences of the helical regions (HR1 and 2) are given in heptad registers of the parent coiled coils on which they were based.

| # | Peptide | Sequence | m/z | |
|---|---|---|---|---|
| | | | Theoretical | Observed |
| **Helical domain 1 (HD1)** | | *gabcdef gabcdef gabcdef gabcdef* | | |
| 1 | HD1-4H | Ac-G EIAAIKE EIAAIKE EIAAIKW EIAAIKE G-$NH_2$ | 3248.810 | 3248.785 |
| 2 | HD1-A@a-4H | Ac-G EAAAIKE EAAAIKE EAAAIKW EAAAIKE G-$NH_2$ | 3080.622 | 3080.651 |
| 3 | HD1-A@a-3H | Ac-G EAAAIKE EAAAIKW EAAAIKE G-$NH_2$ | 2368.247 | 2368.810 |
| 4 | HD1-A@a-2.5H | Ac-G IKE EAAAIKW EAAAIKE G-$NH_2$ | 2025.089 | 2025.722 |
| **Helical domain 2 (HD2)** | | *cdefgab cdefgab cdefgab cdefgab* | | |
| 5 | HD1-4H | Ac-G EIQEQLK EIQEQLK EIQWQLK EIQEQLK G-NH2 | 3704.982 | 3705.882 |
| 6 | HD1-A@a-4H | Ac-G EIQEQAK EIQEQAK EIQWQAK EIQEQAK G-NH2 | 3536.794 | 3537.772 |
| 7 | HD1-A@a-3H | Ac-G EIQEQAK EIQWQAK EIQEQAK G-NH2 | 2710.376 | 2710.527 |
| 8 | HD1-A@a-2.5H | Ac-G QAK EIQWQAK EIQEQAK G-NH2 | 2211.148 | 2211.857 |
| **Linker** | | | | |
| 9 | Linker | Ac-ASPEP QPKPS GDPQS KQTPE PSRSQ-NH2 | 2701.314 | 2701.378 |
| 10 | HD1-L-HD2 | Ac-G IKE EAAAIKW EAAAIKE G-<br>-ASPEP QPKPS GDPQS KQTPE PSRSQ-<br>-G QAK EIQWQAK EIQEQAK G-NH2 | 6821.46 | 6821.234 |

**Supplementary Table S2: Chemically synthesised peptides.** Sequences of the synthesized peptides and the main peaks obtained by MALDI-TOF mass spectrometry. The heptad register is indicated above the sequences (for helical domains 1 and 2). All sequences contained *N*- and *C*-terminal glycine residues (not included in the heptad register assignment) and were *N*-terminally acylated and *C*-terminally amidated.

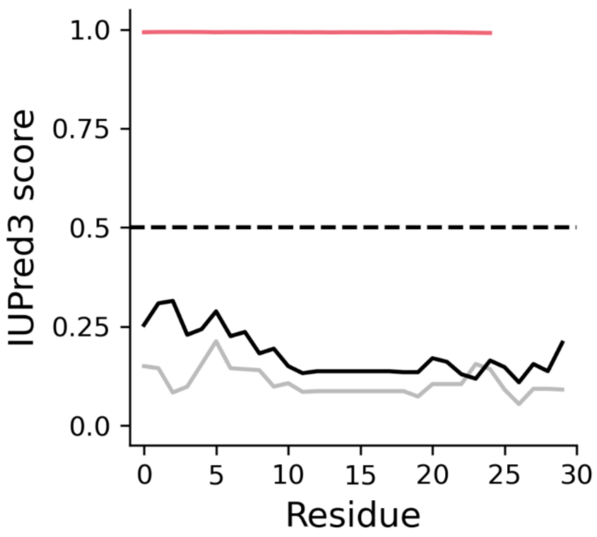


**Supplementary Fig S1: Disorder prediction of the designed HERD-0 components by IUPred3.** Probability of each of the 3 designed components of HERD-0 being part of a disordered region is given by a value between 0 and 1, with a cut-off of 0.5 given by a hashed black line. Pink, linker; grey, helical region 1; solid black. helical region 2. IUPred3 predictions were generated using long disorder parameters with no smoothing applied.

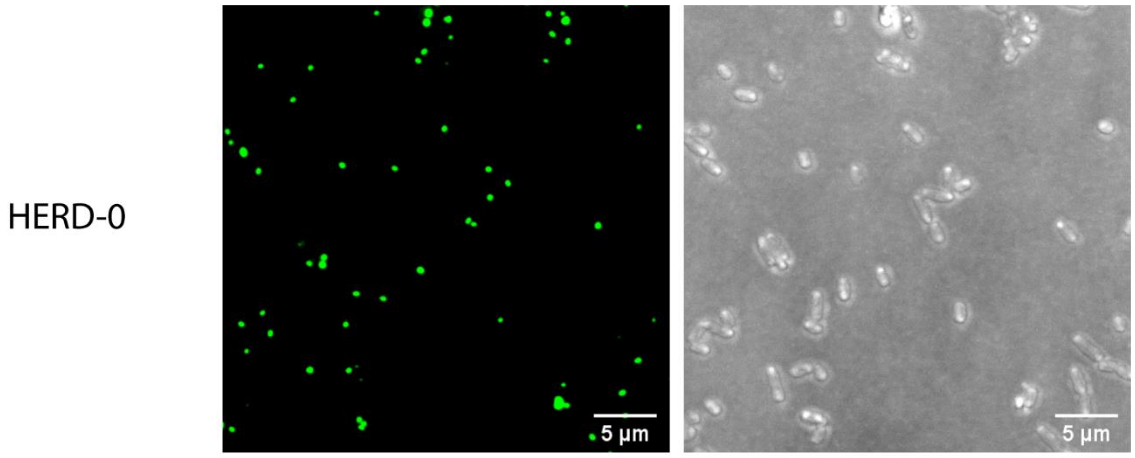


**Supplementary Fig S2: Confocal microscopy images of HERD-0-GFP expressed in *E. coli*.** mEmerald fluorescence at 488 nm (green) and brightfield transmission (grey).

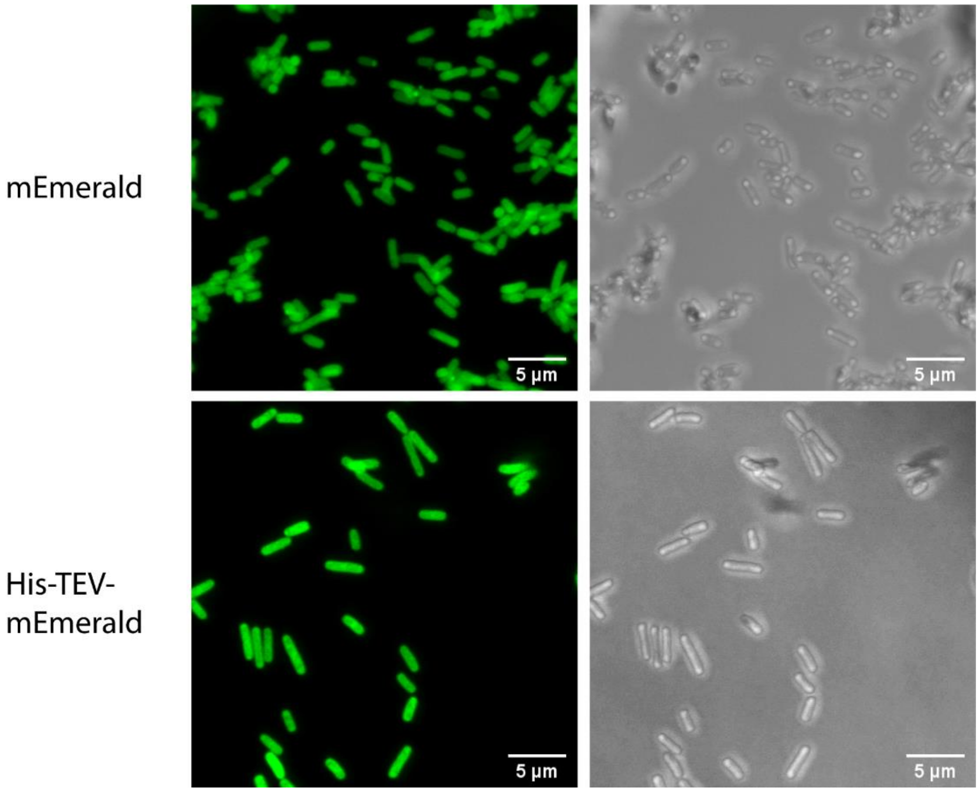


**Supplementary Fig S3: Confocal microscopy images of mEmerald and His-TEV-mEmerald expressed in *E. coli*.** mEmerald fluorescence at 488 nm (green) and brightfield transmission (grey).

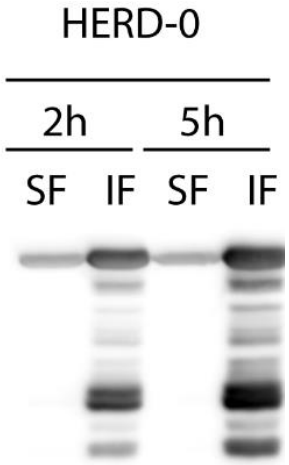


**Supplementary Fig S4: Western blot of HERD-0-GFP from *E. coli* cell lysates.** Cell lysates 2 hours (2h) and 5 hours (5h) after induction were separated into their soluble fractions (SF) and insoluble fractions (IF) before analysis by SDS-PAGE and western blotting against the *N*-terminal His epitope tag

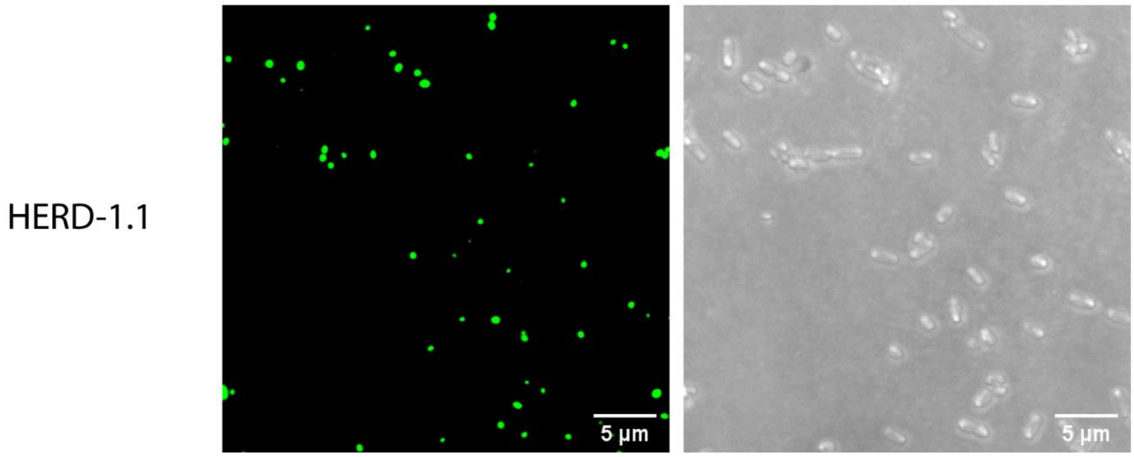


**Supplementary Fig S5: Confocal microscopy images of HERD-1.1 expressed in *E. coli*.** mEmerald fluorescence at 488 nm (green) and brightfield transmission (grey).

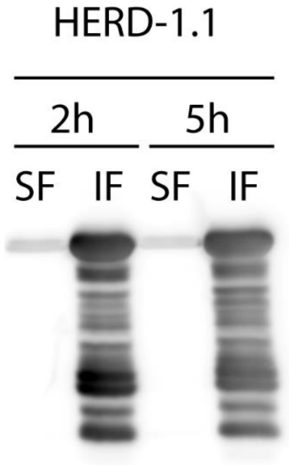


**Supplementary Fig S6: Western blot of HERD-1.1-GFP from *E. coli* cell lysates.** Cell lysates 2 hours (2h) and 5 hours (5h) after induction were separated into their soluble fractions (SF) and insoluble fractions (IF) before analysis by SDS-PAGE and western blotting against the *N*-terminal His epitope tag

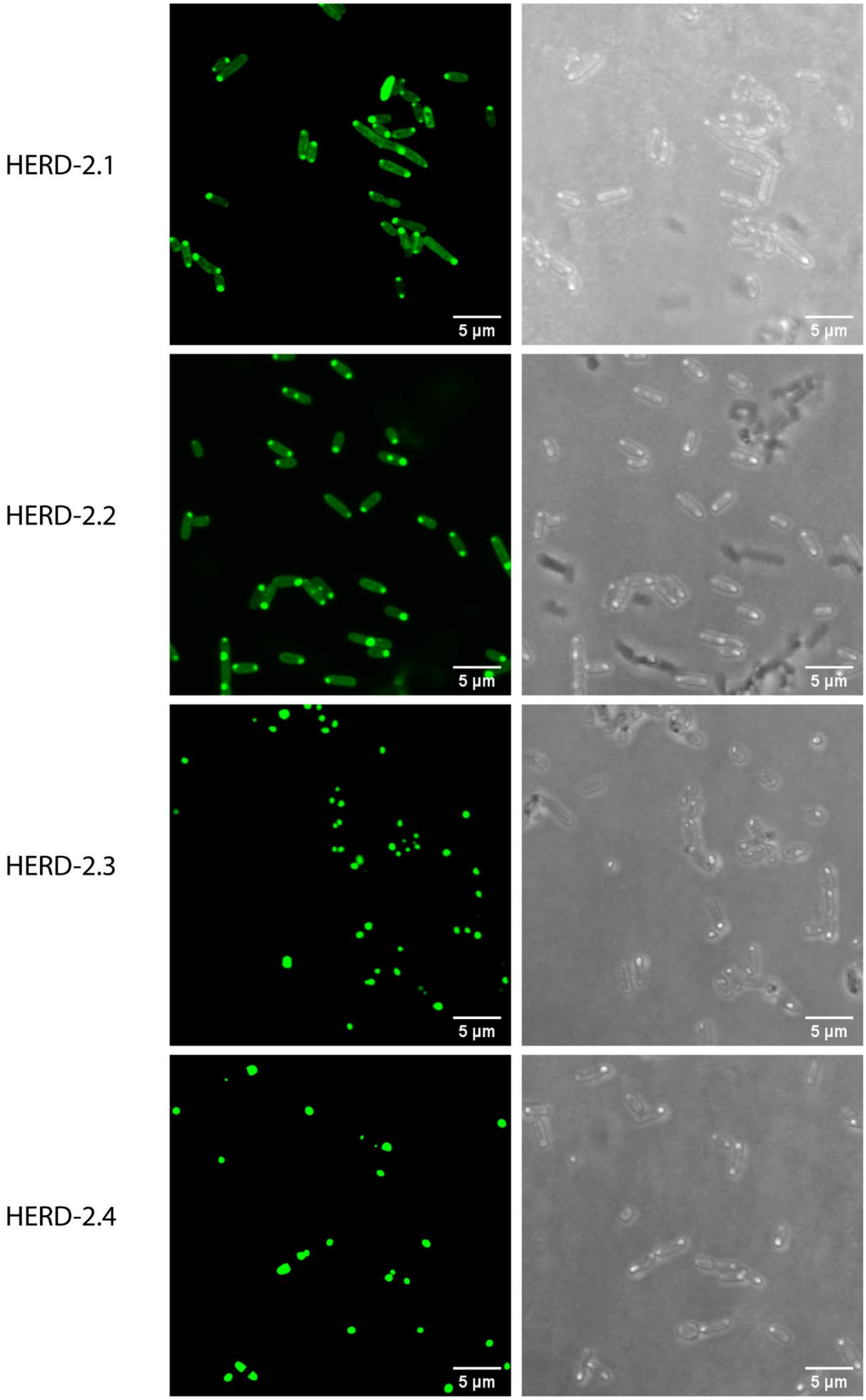


**Supplementary Fig S7: Confocal microscopy images of HERD-2.1 through HERD-2.4 expressed in *E. coli*.** mEmerald fluorescence at 488 nm (green) and brightfield transmission (grey).

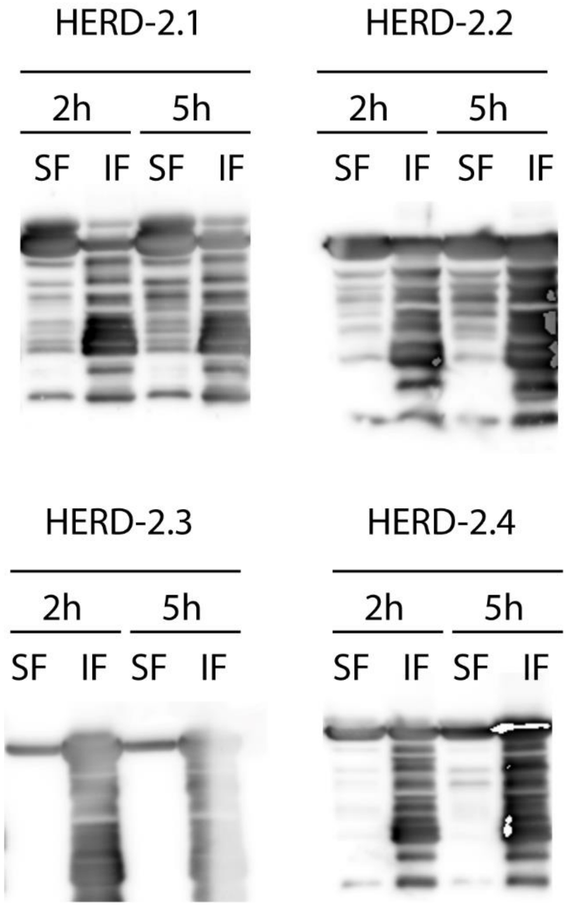


**Supplementary Fig S8: Western blots of HERD-2.1 through HERD-2.4 fusions from *E. coli* cell lysates.** Cell lysates 2 hours (2h) and 5 hours (5h) after induction were separated into their soluble fractions (SF) and insoluble fractions (IF) before analysis by SDS-PAGE and western blotting against the *N*-terminal His epitope tag.

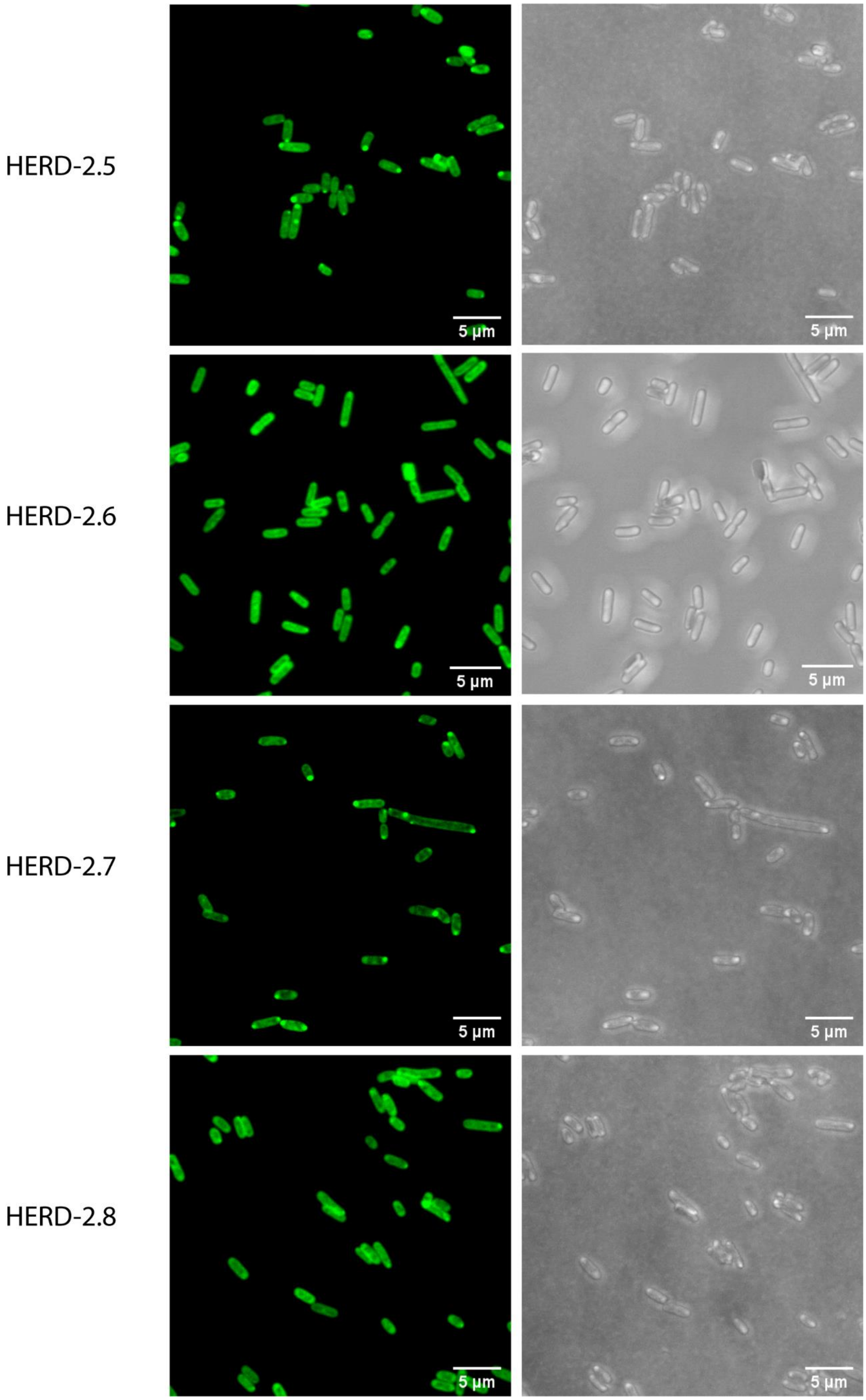


**Supplementary Fig S9: Confocal microscopy images of HERD-2.5 through HERD-2.8 expressed in *E. coli*.** mEmerald fluorescence at 488 nm (green) and brightfield transmission (grey).

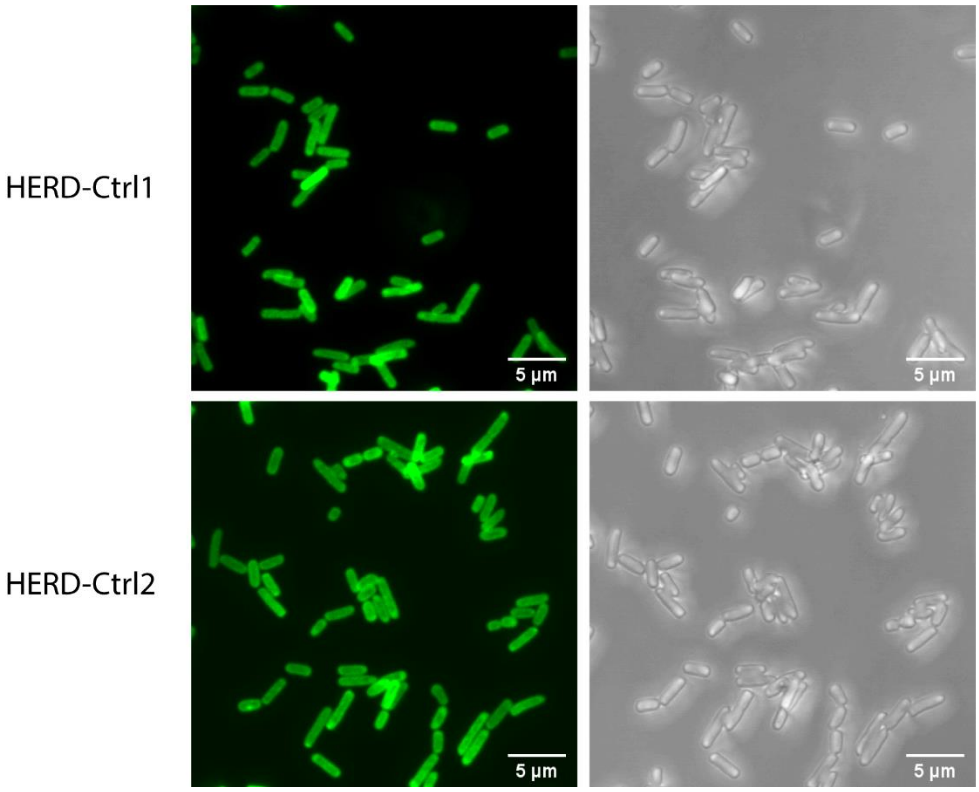


**Figure S10: Confocal microscopy images of HERD-Ctrl1 and HERD-Ctrl2 expressed in *E. coli*.** mEmerald fluorescence at 488 nm (green) and brightfield transmission (grey).

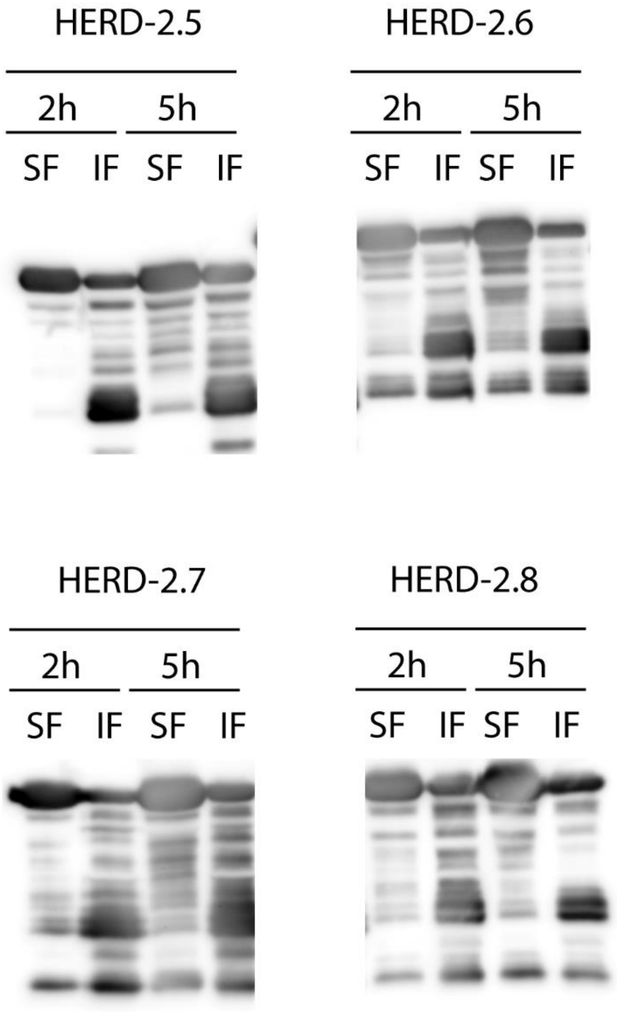


**Supplementary Fig S11: Western blots of HERD-2.5 through HERD-2.8 fusions from *E. coli* cell lysates.** Cell lysates 2 hours (2h) and 5 hours (5h) after induction were separated into their soluble fractions (SF) and insoluble fractions (IF) before analysis by SDS-PAGE and western blotting against the *N*-terminal His epitope tag.

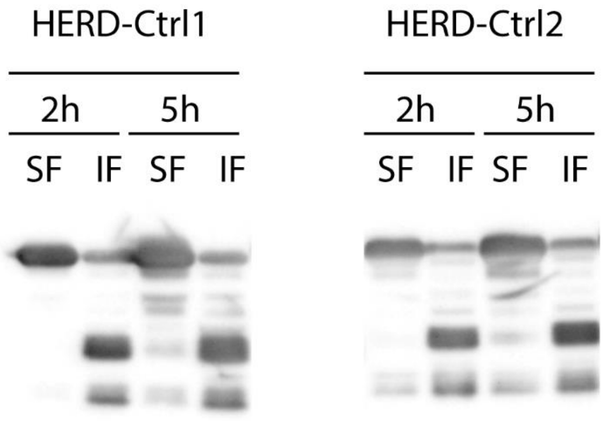


**Supplementary Fig S12: Western blots of HERD-Ctrl1 and HERD-Ctrl2 fusions from *E. coli* cell lysates.** Cell lysates 2 hours (2h) and 5 hours (5h) after induction were separated into their soluble fractions (SF) and insoluble fractions (IF) before analysis by SDS-PAGE and western blotting against the *N*-terminal His epitope tag.

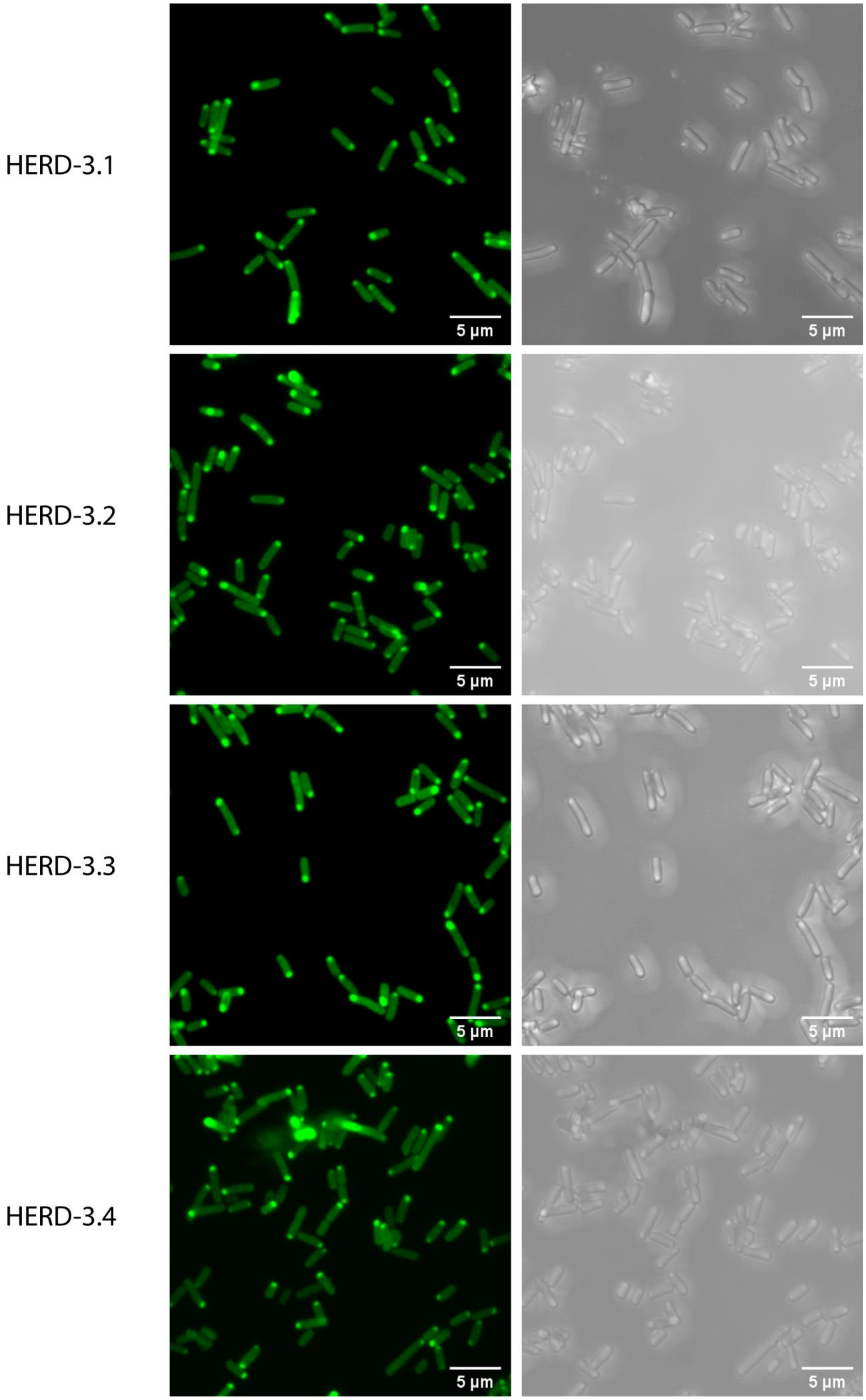


**Supplementary Fig S12: Confocal microscopy images of HERD-3.1 through HERD-3.4 expressed in *E. coli*.** mEmerald fluorescence at 488 nm (green) and brightfield transmission (grey).

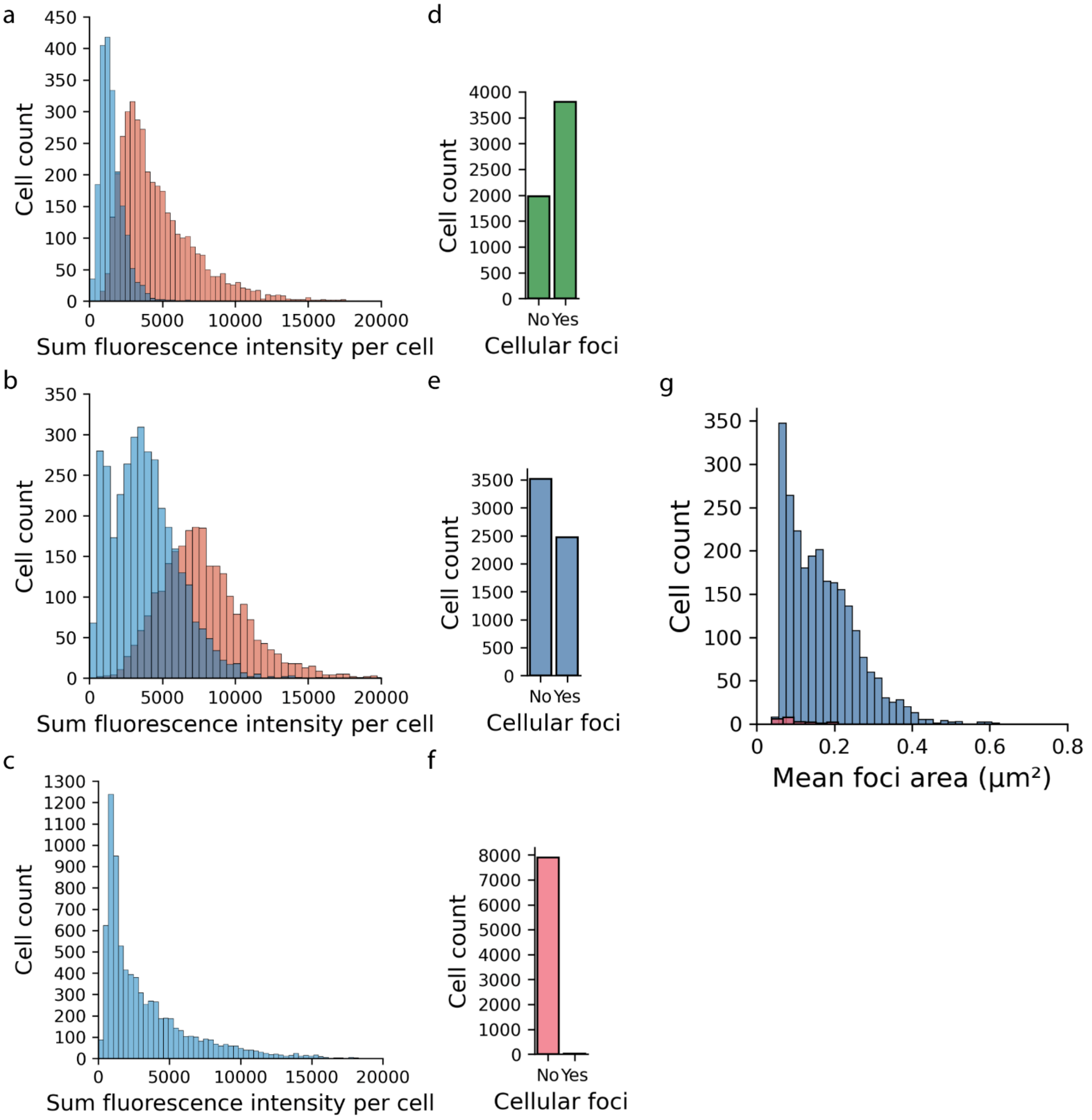


**Supplementary Fig S13: Automated image analysis of HERD–GFP fusions from confocal microscopy. a, b & c,** Histogram of cells expressing HERD-0–GFP **(a)**, HERD-2.2–GFP **(b)**, and HERD-Ctrl2–GFP **(c).** Images for analysis taken across 6 hours of protein expression to gather variable intracellular protein concentrations. Cells detected as containing no protein condensates are labelled blue, and cells containing at least one protein condensate are labelled red. **d, e, & f,** Histogram of cells expressing HERD-0–GFP **(d)**, HERD-2.2–GFP **(e)**, and HERD-Ctrl2–GFP **(f)** analysed for the presence of protein condensates. **g,** Histogram of mean protein condensate area in cells expressing HERD-2.2–GFP (blue) and HERD-Ctrl2–GFP (pink).

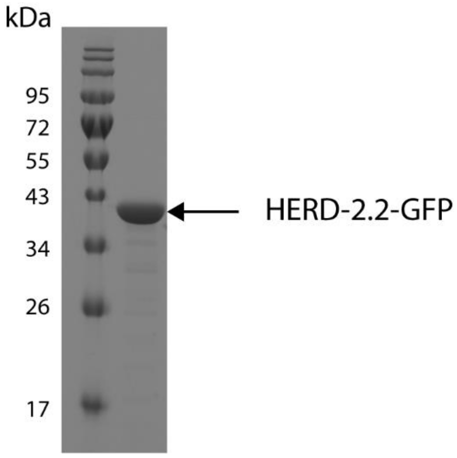


**Supplementary Fig S14: SDS-PAGE gel of purified HERD-2.2–GFP.** Molecular weight marker is NEB colour pre-stained protein standard broad range (10 – 250 kDa).

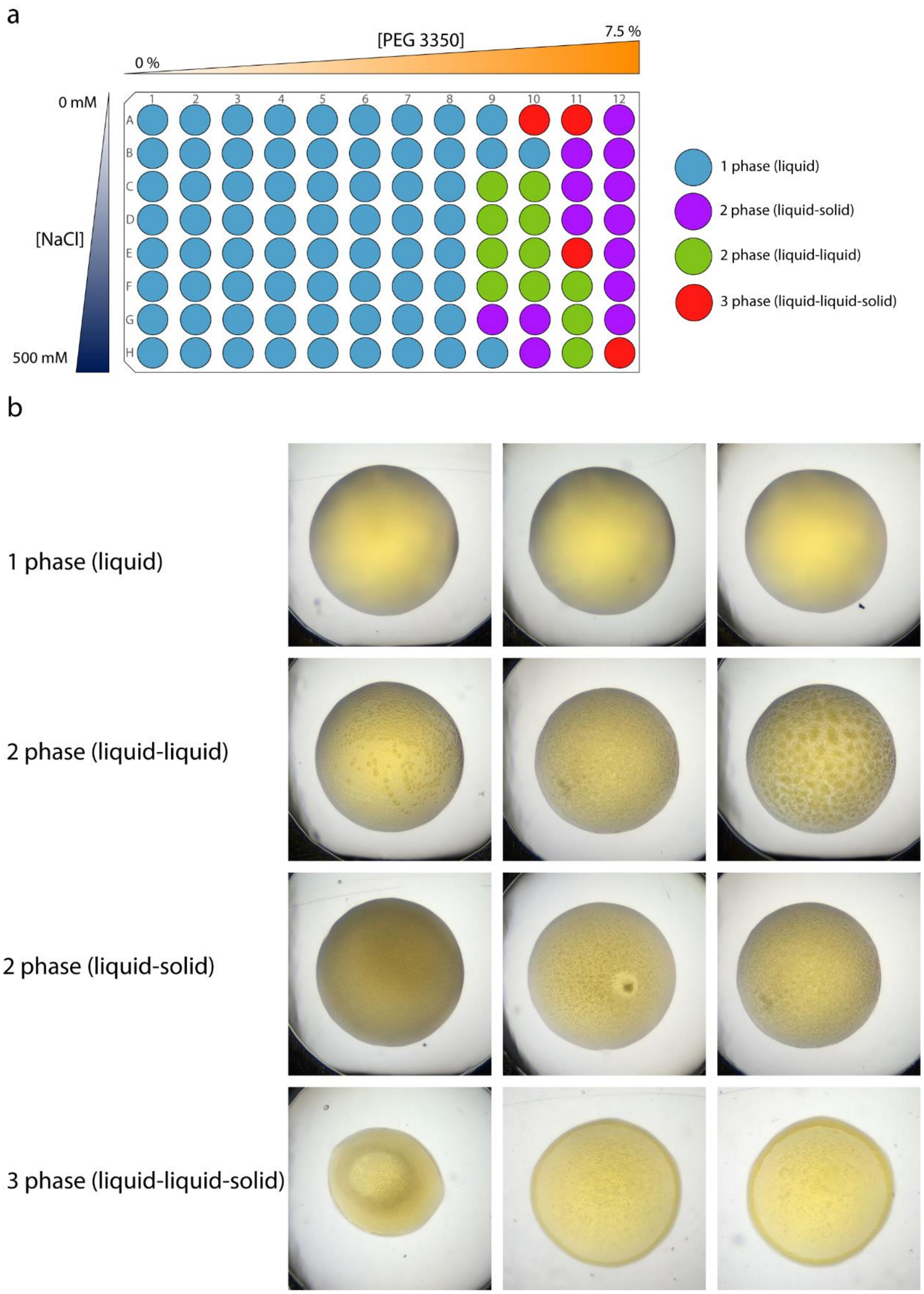


**Supplementary Fig S15: Image based buffer screen for phase separation of HERD-2.2–GFP. a,** Tabulated buffer screen for phase separation of HERD-2.2–GFP. **b,** Representative images of the properties described in the tabulated screen for phase separation. Classification was made on the following observed properties: 1 phase was classified based on the absence of any protein precipitation or de-mixing. 2 phases (liquid-liquid) was classified based on the appearance of de-mixed particles that appeared largely spherical and had smooth boundaries between the two phases. 2 phases (liquid-solid) was classified based on the appearance off granular particles that were not spherical and had coarse or granular boundaries between the 2 phases, and darkening of the solution. 3 phases (liquid-liquid-solid) was classified based on the appearance of both morphologies within the same droplet. Conditions – Varying from 0 – 500 mM NaCl, 0 – 7.5% PEG 3350, with 20 mM Tris-HCl pH 7.5, 400 µM HERD-2.2–GFP. Total volume 0.8 µl.

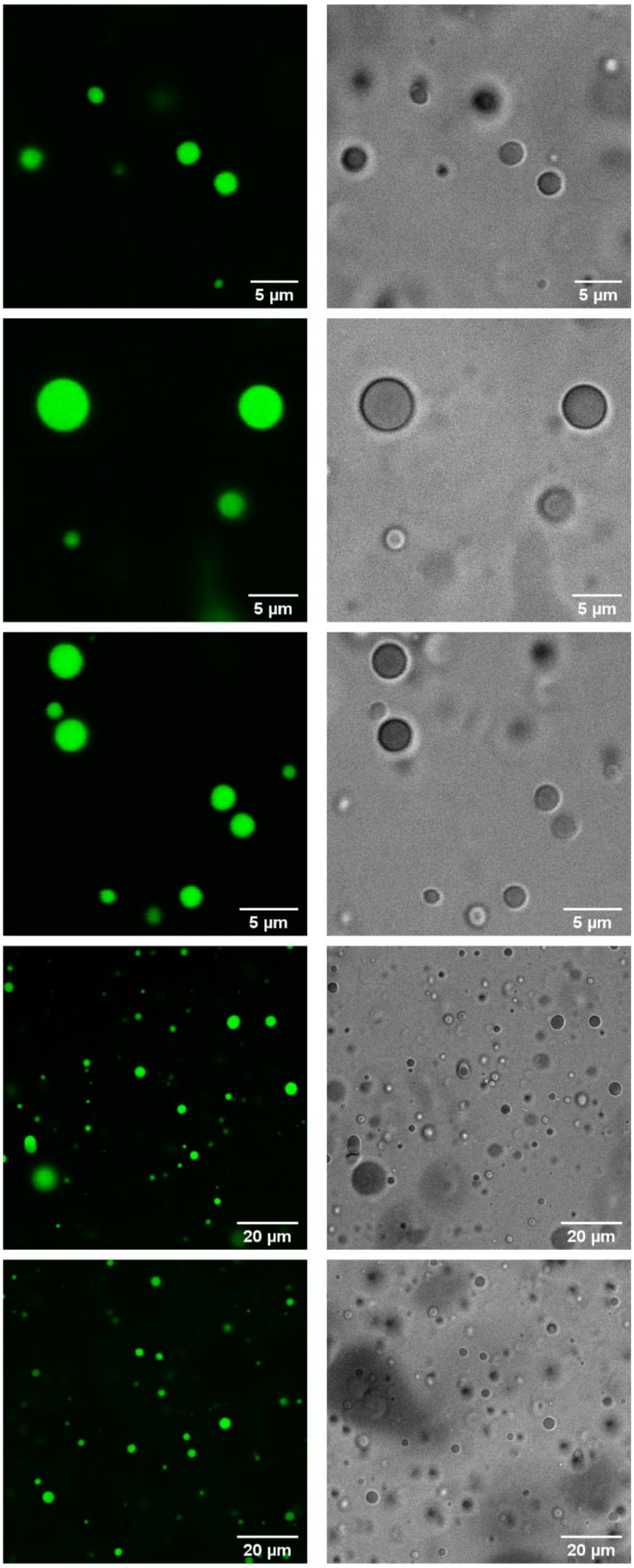


**Supplementary Fig S16: Confocal microscopy images of HERD-2.2-GFP droplets *in vitro*.** mEmerald fluorescence at 488 nm (green) and brightfield transmission (grey). Conditions: 125 mM NaCl, 4% PEG 3350, 20 mM Tris-HCl pH 7.5, 1 mM HERD-2.2–GFP.

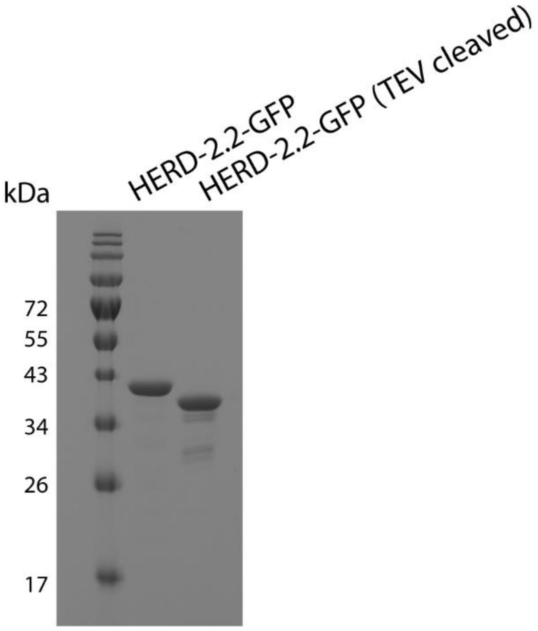


**Supplementary Fig S17: SDS-PAGE of HERD-2.2-GFP before and after TEV cleavage.** Molecular weight marker is NEB colour pre-stained protein standard broad range (10 – 250 kDa).

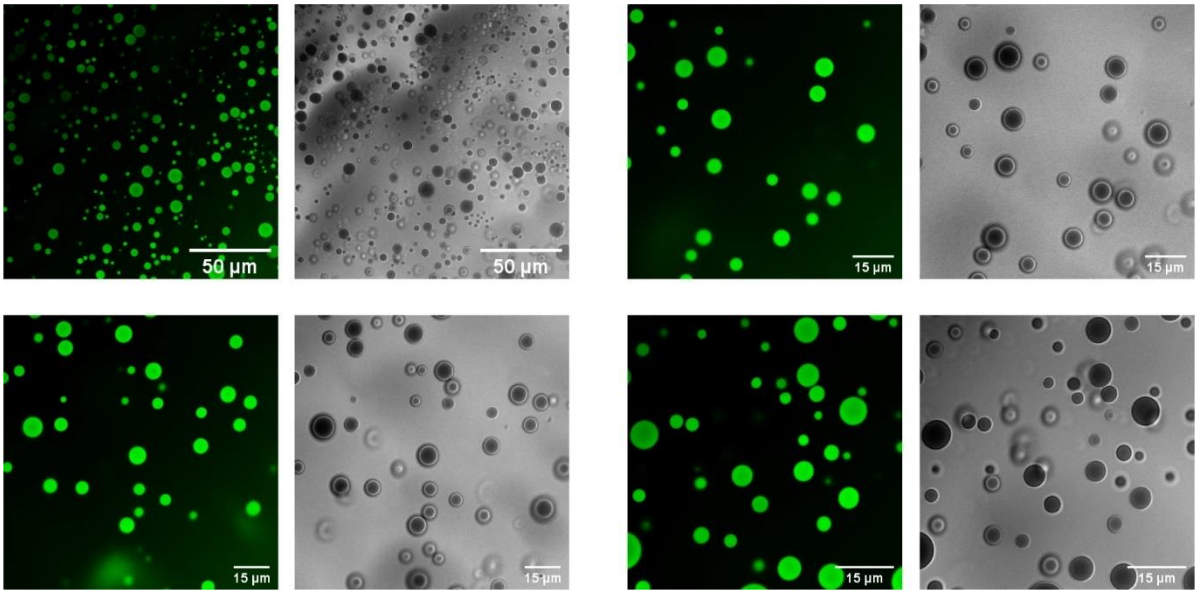


**Supplementary Fig S18: Confocal microscopy images of droplets *in vitro* formed by HERD-2.2-GFP after TEV cleavage.** mEmerald fluorescence at 488 nm (green) and brightfield transmission (grey). Conditions: 125 mM NaCl, 10 % PEG 3350, 20 mM Tris-HCl pH 7.5, 2 mM HERD-2.2–GFP (cleaved).

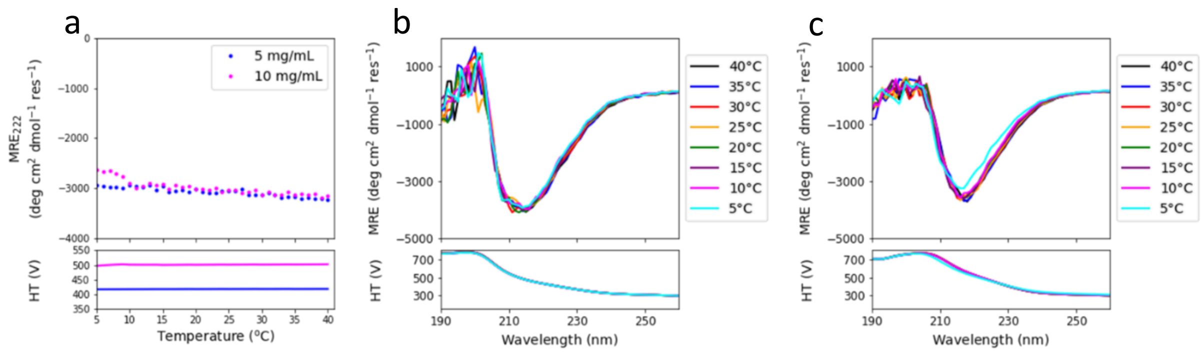


**Supplementary Fig S19: Circular dichroism spectroscopy of HERD-2.2-GFP at varying temperature and concentration**. **a,** Changes in $MRE_{222}$ at 5 and 10 mg/ml protein concentration. **b & c,** CD spectra for 5 and 10 mg/ml protein corresponding to the curves on panel **a**. Spectra were recorded from 40 – 5 °C with a 5 °C step. Conditions: 20 mM Tris-HCl pH 7.5, 125 mM NaCl, 4% PEG3350, 0.1 mm path length,.

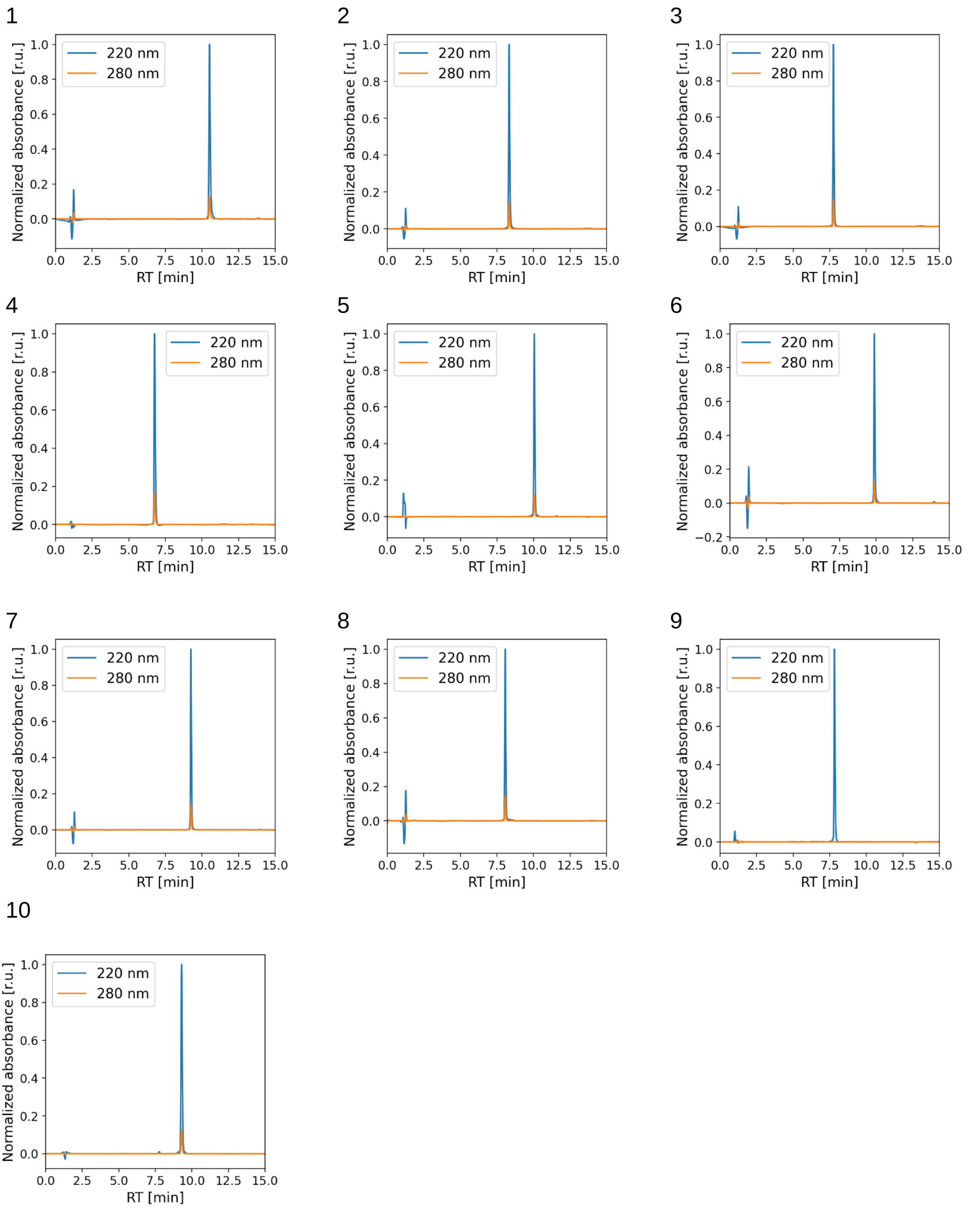


**Supplementary Fig S20: Representative analytical HPLC traces of synthesised peptides 1-10 monitoring absorbance at 220 nm and 280 nm.** Absorbances are normalised intensities to the maxima at 220 nm. Peptides are numbered as in Supplementary Table S2. HPLC was performed using a C18 reverse phase column (Phenomenex Kinetex, 5 µm particle size, 100 Å pore size, 100 × 4.6 mm), a gradient of Buffer A (0.1 % TFA in $H_2O$) and Buffer B (0.1 % TFA in MeCN), at a flow rate of between 1-3 ml/min.

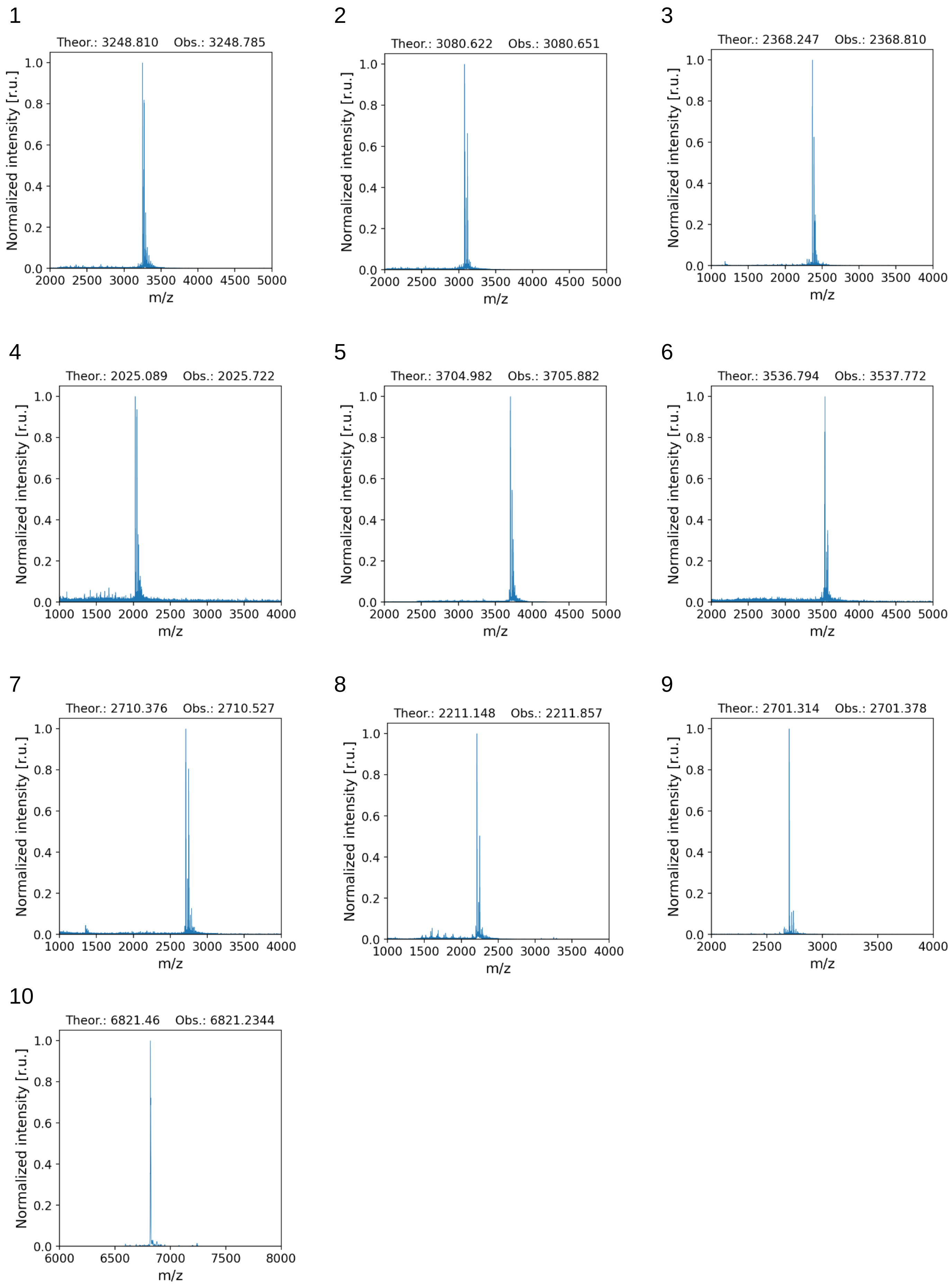


**Supplementary Fig S21. Representative MALDI-TOF spectra for the chemically synthesised peptides 1-10.** Mass spectra of the synthetic peptides given in Supplementary Table S2.

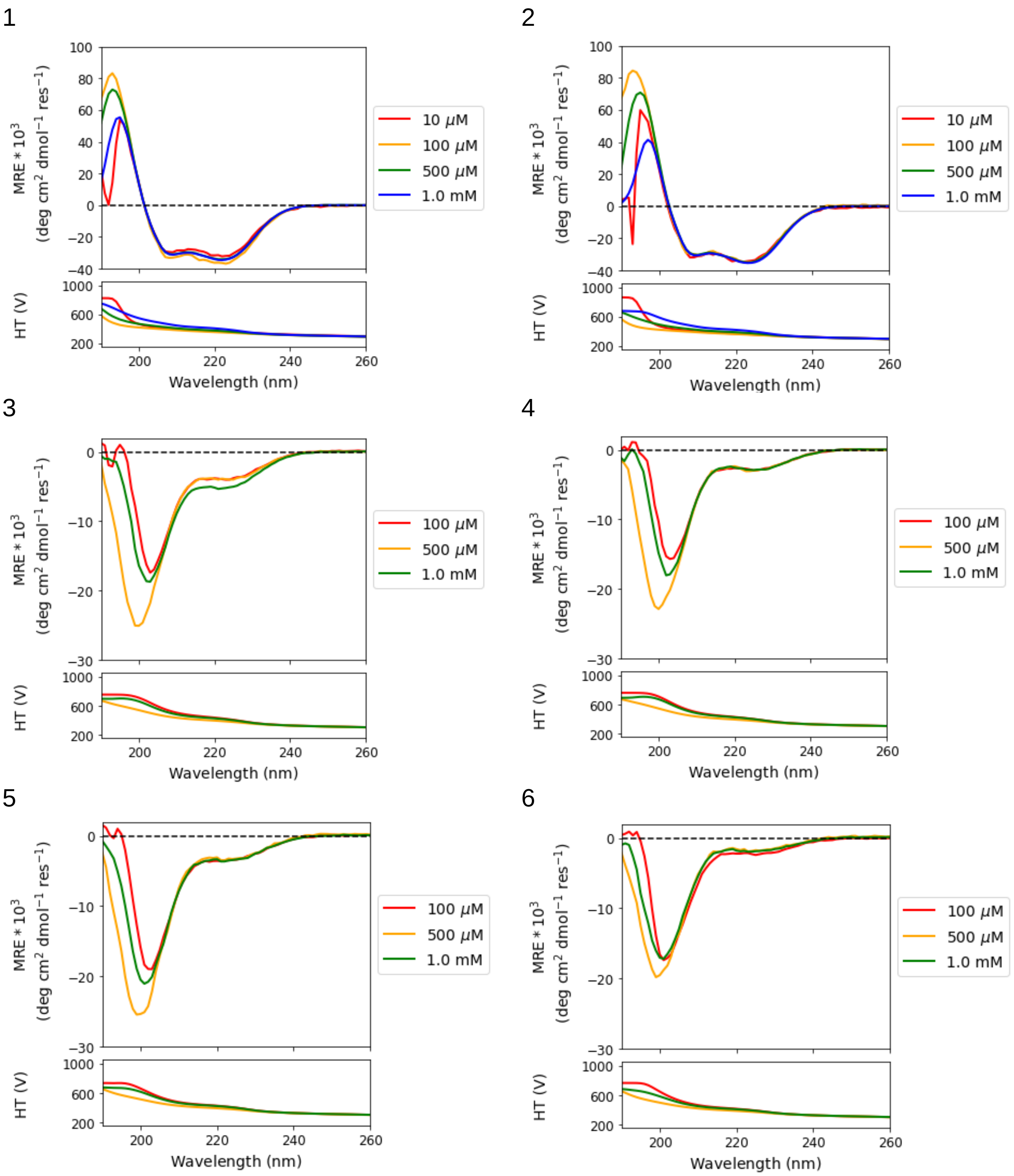
1
2
3
4
5
6
MRE * 10$^3$ (deg cm$^2$ dmol$^{-1}$ res$^{-1}$)
HT (V)
Wavelength (nm)
10 μM
100 μM
500 μM
1.0 mM

7

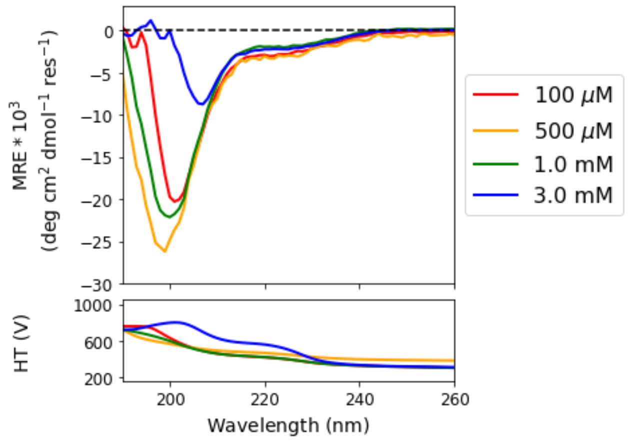


8

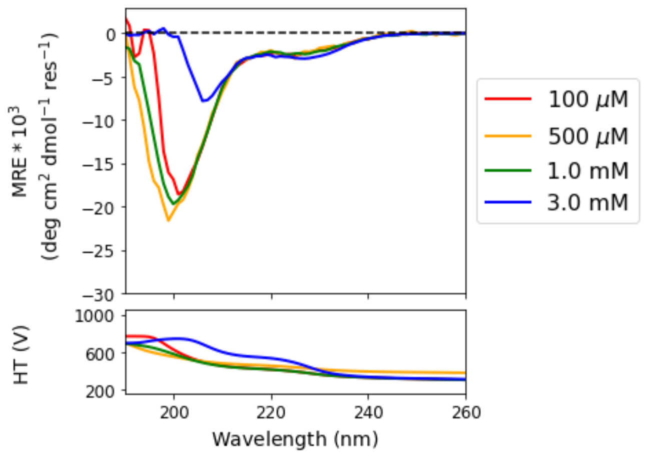


9

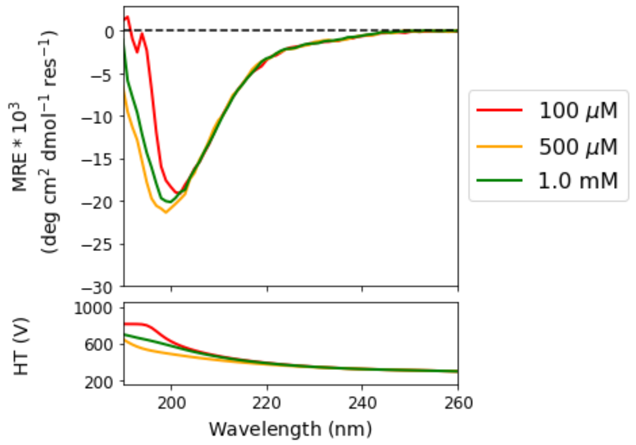


10

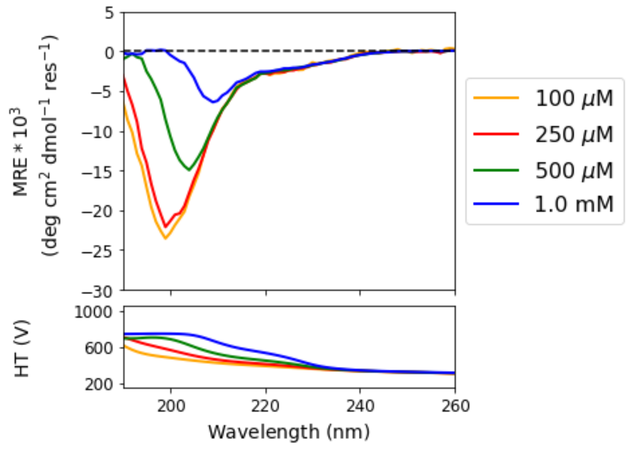


**Supplementary Fig S22. Circular dichroism spectra for the chemically synthesised peptides 1 – 10.** Peptide sequences are given in Supplementary Table S2. Conditions: 20 mM Tris-HCl pH 7.5, 125 mM NaCl, 5 °C, with varied peptide concentrations.

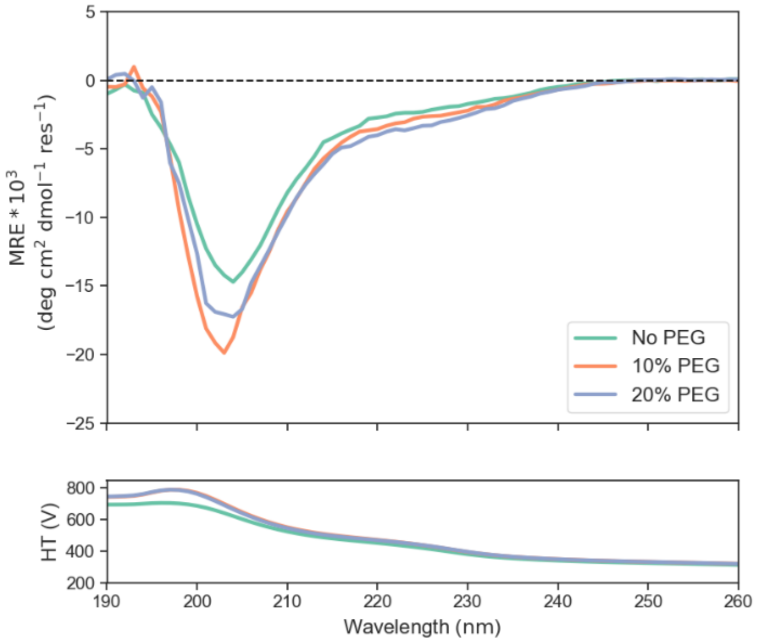


**Supplementary Fig S23: CD spectra for the peptide 10 at 500 μM in the presence of different PEG 3350 concentrations.** Conditions: 20 mM Tris-HCl pH 7.5, 125 mM NaCl, 5 °C.

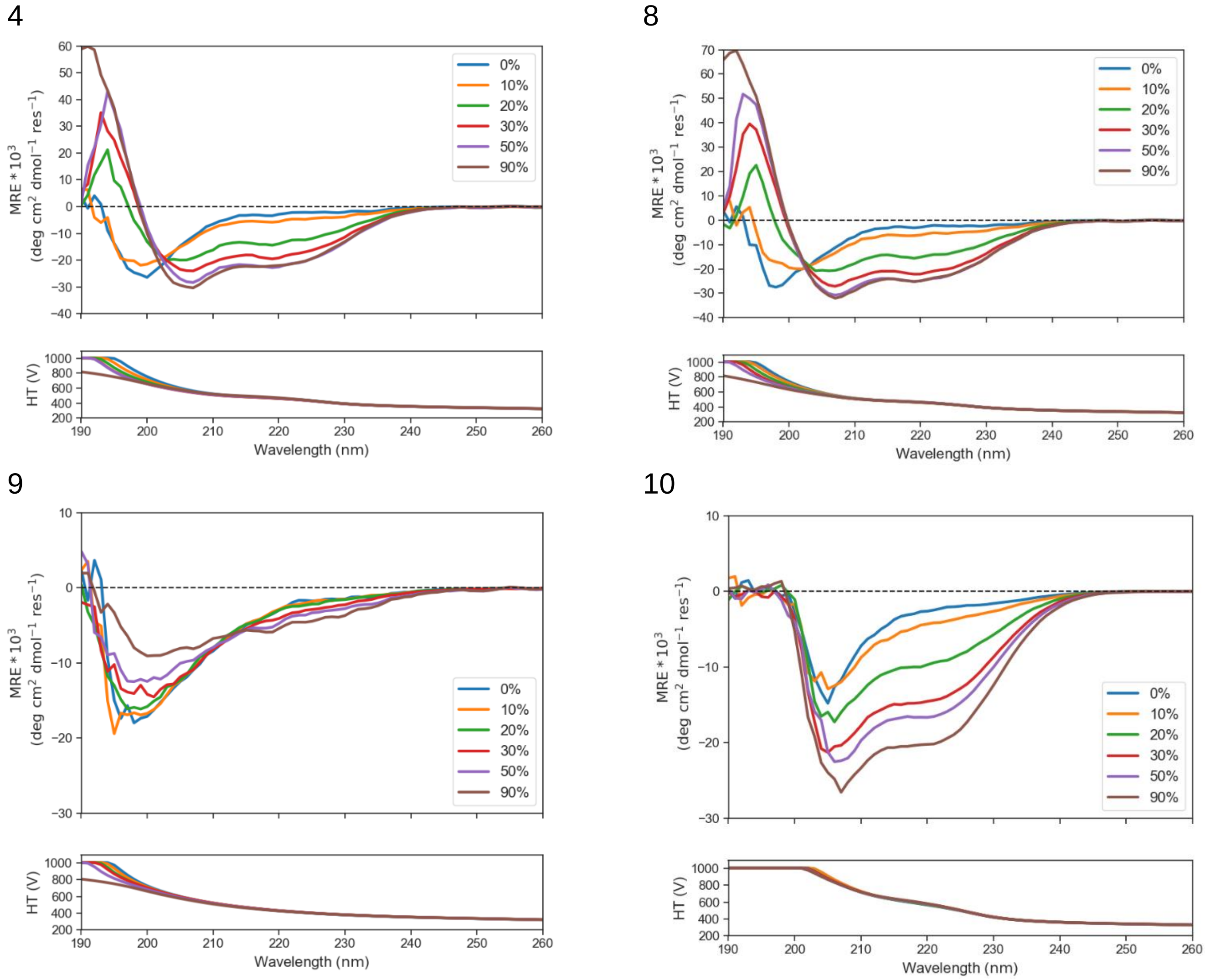


**Supplementary Fig S24: CD spectra for the peptides 4 and 8 – 10 at 100 µM in the presence of different concentrations of TFE (0 – 90%).** Peptide sequences are given in Supplementary Table S2. Conditions: 20 mM Tris-HCl pH 7.5, 125 mM NaCl, 5 °C.

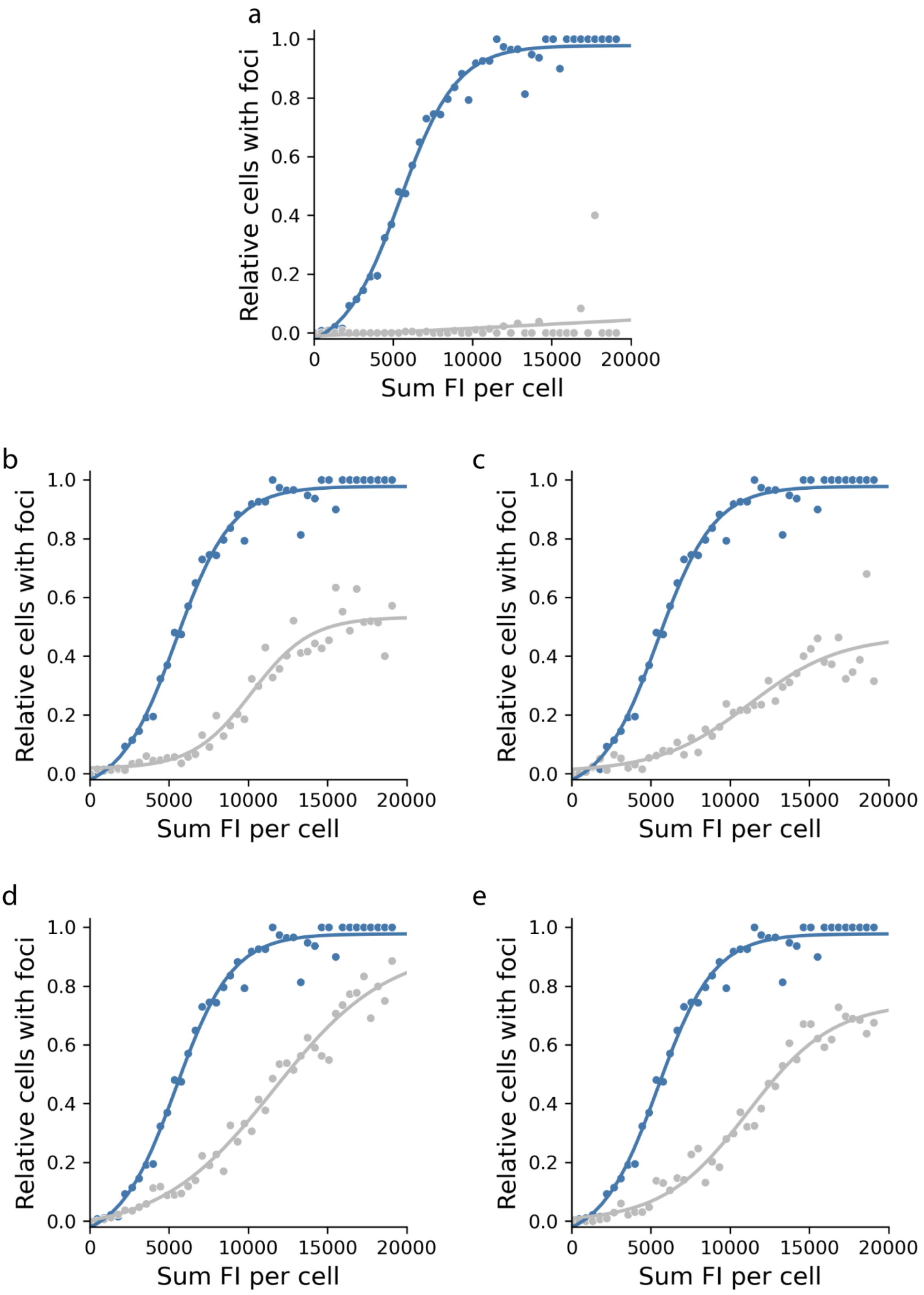


**Supplementary Fig S25: Automated image analysis of helix destabilising HERD-GFP constructs.** Comparison of protein condensate formation between cells expressing HERD-2.2–GFP (blue, n = 5993) and cells expressing constructs with the destabilising mutations (grey). Helix disrupted HERD fusions in each panel: **a**, HERD-Ctrl3–GFP, n = 4141; **b**, HERD-Ctrl4–GFP, n = 5530; **c**, HERD-Ctrl5–GFP, n = 5554; **d**, HERD-Ctrl6–GFP, n = 4339; **e** – HERD-Ctrl7–GFP, n = 4113.

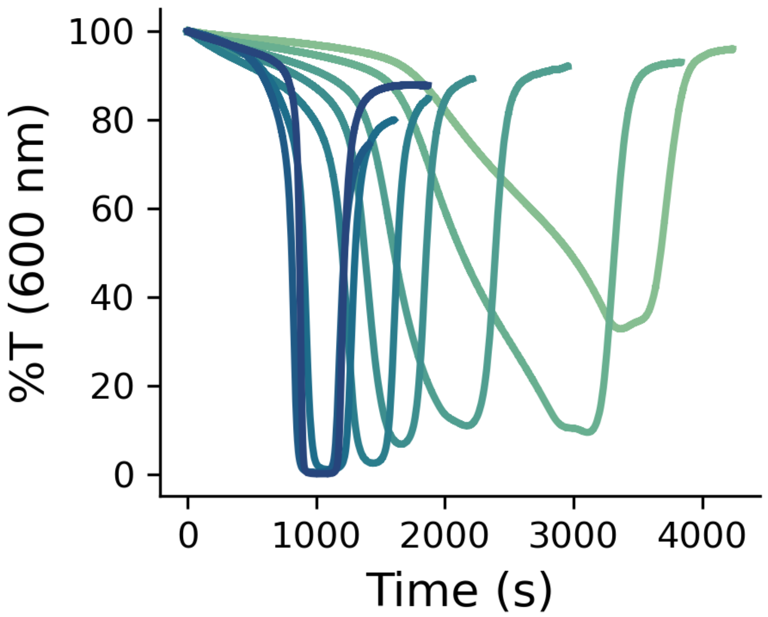


**Supplementary Fig S26: Cloud-point measurements for determining the binodal phase boundary of HERD-2.2–GFP *in vitro* with respect to temperature.** Phase separation was measured by changes in transmission (%T, 600 nm) over time with HERD-2.2–GFP varied from 72 µM to 1 mM. Conditions: 20 mM Tris-HCl pH 7.5, 125 mM NaCl , 4% PEG 3350.

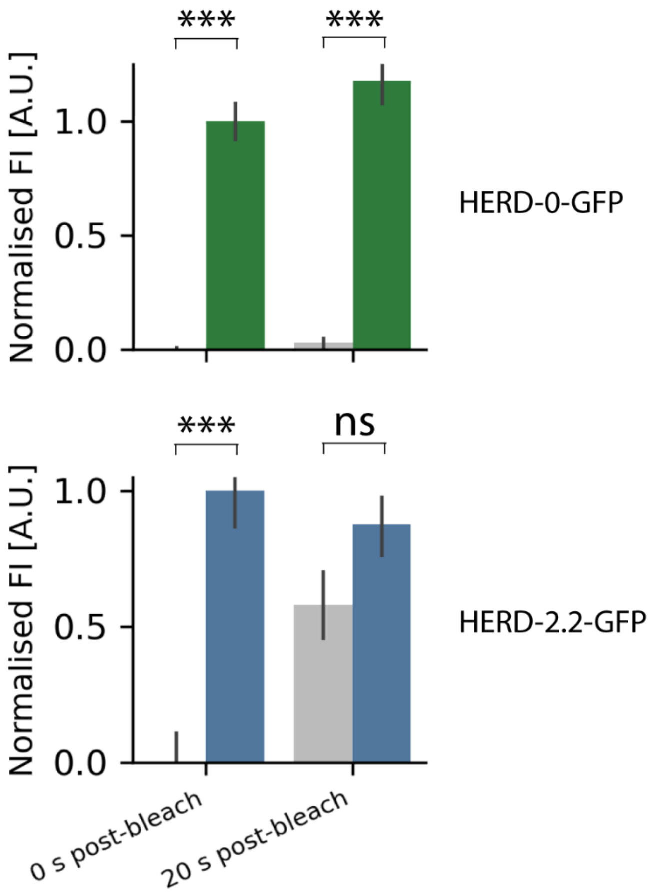


**Supplementary Fig S27: Quantification of asymmetrical bleaching of HERD-0–GFP (green) and HERD-2.2–GFP (blue) condensates in cells**. Normalised fluorescence intensity within the directly bleached area is shown in grey, fluorescence intensity outside of the bleached area within the same droplet is shown in green (HERD-0–GFP; n = 19) and blue (HERD-2.2–GFP; n = 13). Samples are condensates within discrete cells measured independently. Fluorescence intensity was normalised relative to the mean fluorescence intensity in the bleached area (0) and the mean fluorescence intensity outside of the bleached area, in the same droplet (1). Error bars represent the standard error. Statistical testing used a two tailed t-test comparing the normalised fluorescence intensity within (in-spot) the bleached spot and outside (ex-spot) of the bleached spot. Samples were confirmed to conform to assumptions for equal variance and normality by Shapiro testing and Fligner-Killeen testing respectively. P-value < 0.001 for all comparisons aside from HERD-2.2–GFP 20 s post-bleach (p-value = 0.0958, accepted null hypothesis). Mean and standard deviation for all samples as follows. HERD-0–GFP: 0 s post bleach; in-spot mean = 0.000, standard deviation = 0.031; ex-spot mean = 1.000, standard deviation = 0.374. 20 s post bleach; in-spot mean = 0.027, standard deviation = 0.080; ex-spot mean = 1.17, standard deviation = 0.471. HERD-2.2–GFP: 0 s post bleach; in-spot mean = 0.000, standard deviation = 0.395, ex-spot: mean = 1.000, standard deviation = 0.492. 20 s post bleach; in-spot mean = 0.579, standard deviation = 0.444; ex-spot mean = 0.875, standard deviation = 0.391.

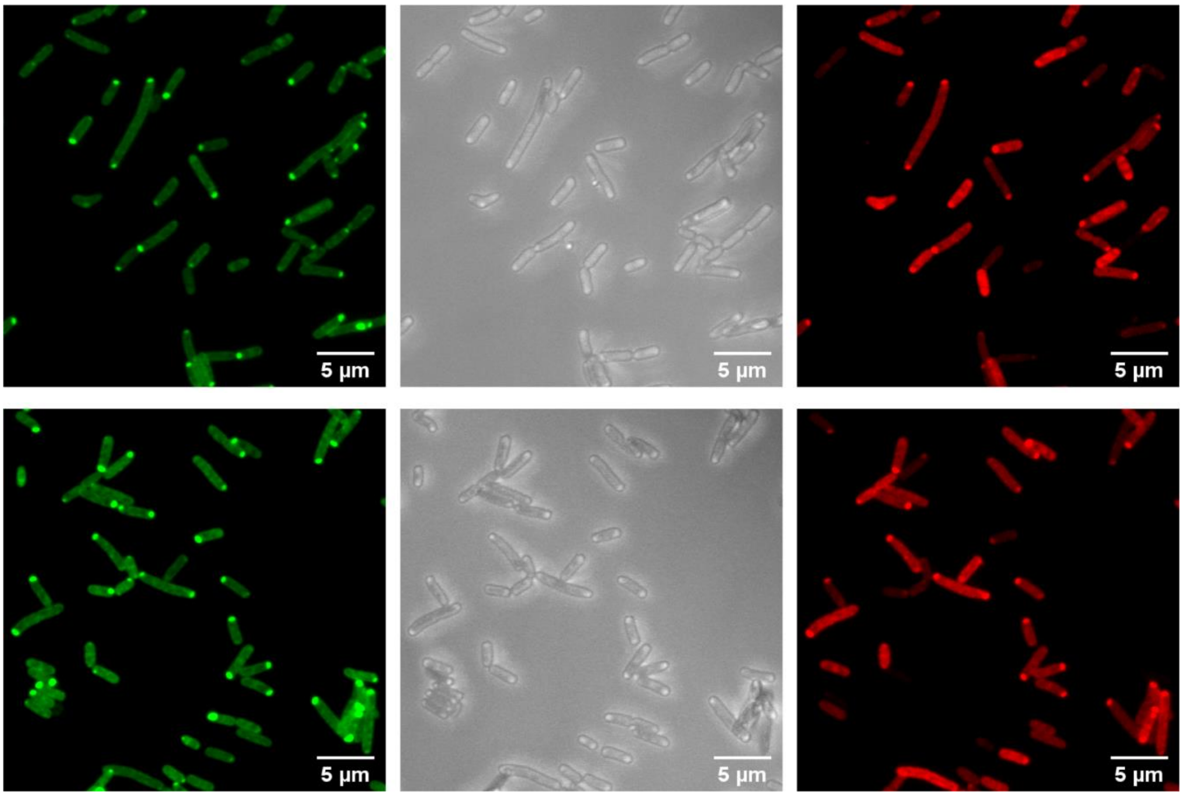


**Supplementary Fig S28: Examples of confocal microscopy images of HERD-2.2–GFP co-expressed with HERD-2.2–mCherry in *E. coli*.** mEmerald fluorescence at 488 nm (green), brightfield transmission (grey), and mCherry fluorescence at 561 nm (red).

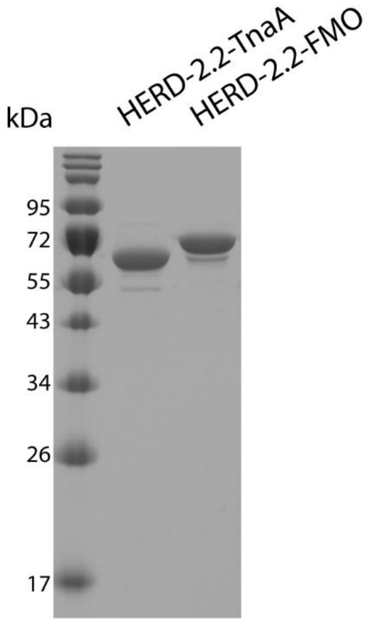


**Supplementary Fig S29: SDS-PAGE of HERD-2.2 fusions with TnaA and FMO.** Molecular weight marker is NEB colour pre-stained protein standard broad range (10 – 250 kDa).

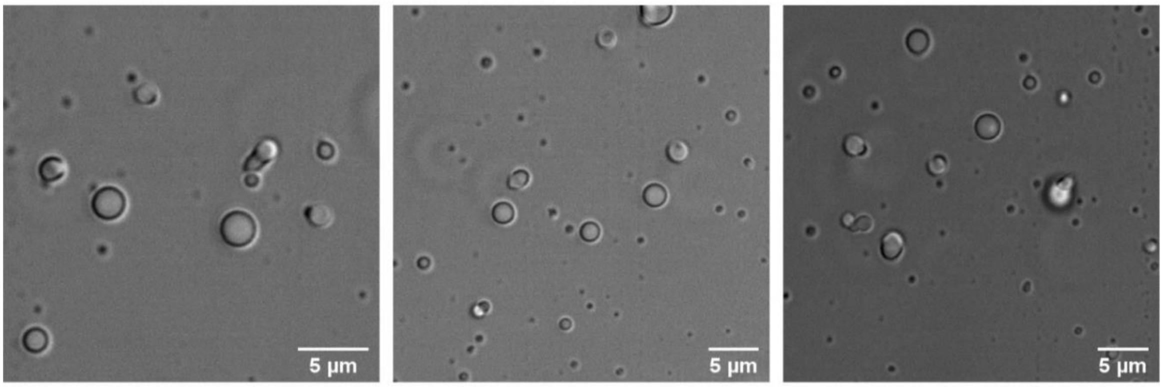


**Supplementary Fig S30: Brightfield transmission images of de-mixed droplets formed by HERD-2.2 fusion with TnaA.** Conditions: 200 mM NaCl, 8% PEG 3350, 40 mM Bis-Tris-HCl pH 6, 120 µM HERD-2.2–TnaA.

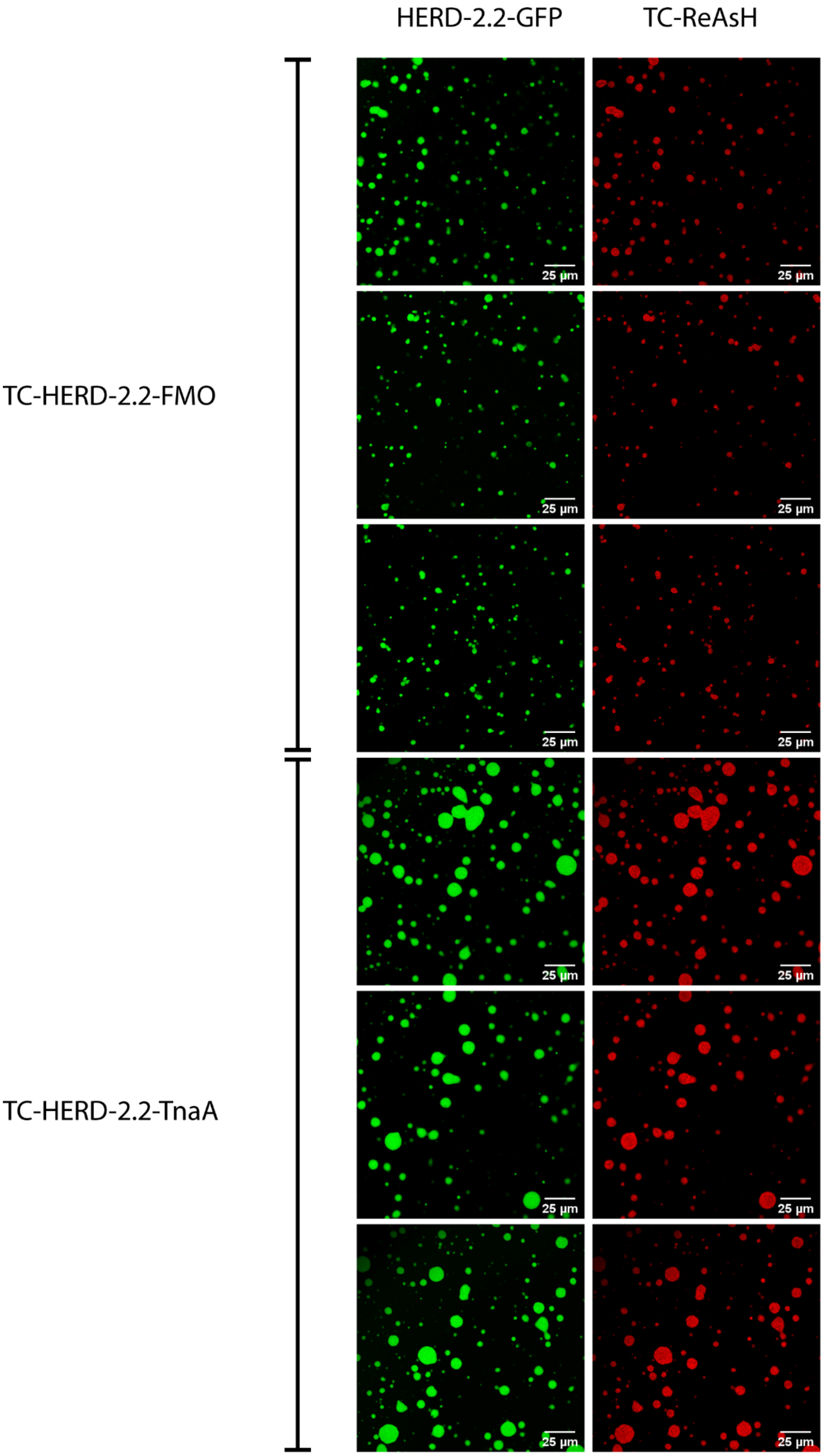


**Supplementary Fig S31: *In vitro* confocal microscopy images of enriched TC tagged HERD-2.2 fusions.** mEmerald fluorescence at 488 nm corresponding to HERD-2.2–GFP (green), and TC-ReAsH fluorescence at 561 nm corresponding to the enriched tetra-cysteine tagged enzyme fusions (red). Conditions: 125 mM NaCl, 4% PEG 3350, 20 mM Tris-HCl pH 7.5, 2 mM HERD-2.2–GFP, 100 µM HERD-2.2–TnaA or HERD-2-2–FMO.

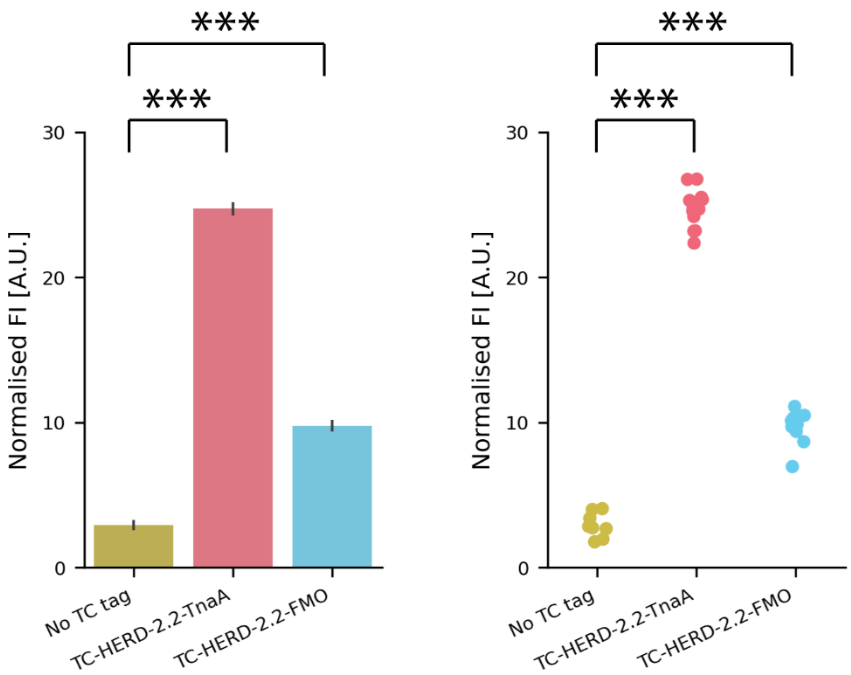


**Supplementary Fig S32: Quantification of enriched TC tagged HERD-2.2 fusions.** Quantification of the fluorescence intensity after excitation at 561 nm (for TC-ReAsH) in de-mixed droplets formed by HERD-2.2–GFP shown as bar chart (left) and scatter plot (right). The points are for discrete droplets measured independently. Statistical analysis was performed by one way ANOVA between the 3 samples followed by TukeyHSD post-hoc test to perform a multiple comparison of means and variance. p-adj < 0.001 for all comparisons. Sample size, mean and standard deviation for each sample as follows. No TC tag: n = 8; mean = 2.92; standard deviation = 0.79. TC-HERD-2.2–TnaA: n = 12; mean = 24.70; standard deviation = 1.30. TC-HERD-2.2–FMO: n = 12; mean = 9.76; standard deviation = 1.03.

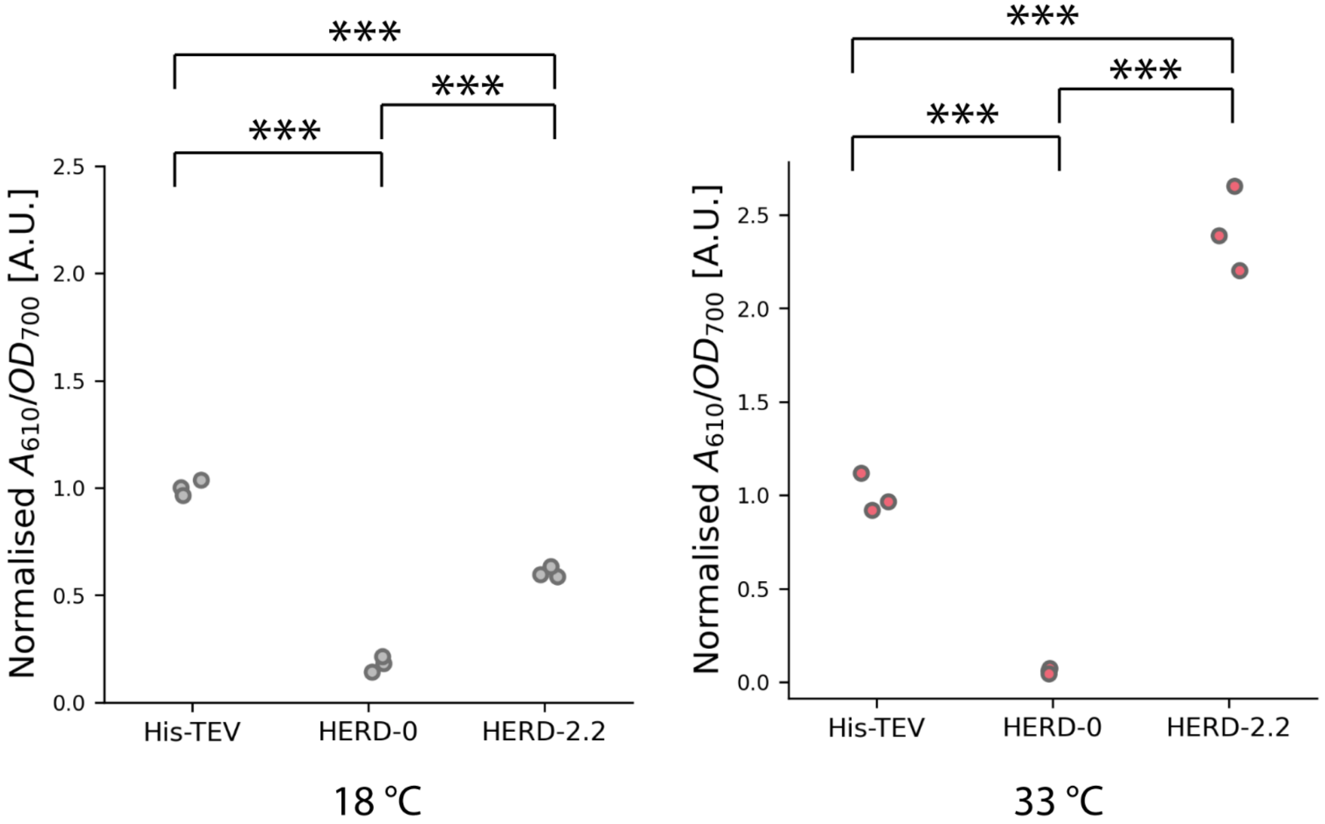


**Supplementary Fig S33: Normalised indigo production in cells co-expressing fusions of TnaA and FMO with complementary GFP-labelled scaffold proteins at 18 °C (grey) and 33 °C (pink).** n=3 for each condition. Condition replicates are discrete cultures measured independently. Statistical analysis was performed by one way ANOVA between the 3 conditions of each experimental temperature, followed by TukeyHSD post-hoc test to perform a multiple comparison of means and variance. p-adj <0.001 for all comparisons. Mean and standard deviation for each condition as follows. 18 °C: His-TEV; mean = 1.000, standard deviation = 0.029; HERD-0; mean = 0.178, standard deviation = 0.030; HERD-2.2; mean = 0.604, standard deviation = 0.020. 33 °C: His-TEV; mean = 1.000, standard deviation = 0.084; HERD-0; mean = 0.0577, standard deviation = 0.012; HERD-2.2; mean = 2.414, standard deviation = 0.185.

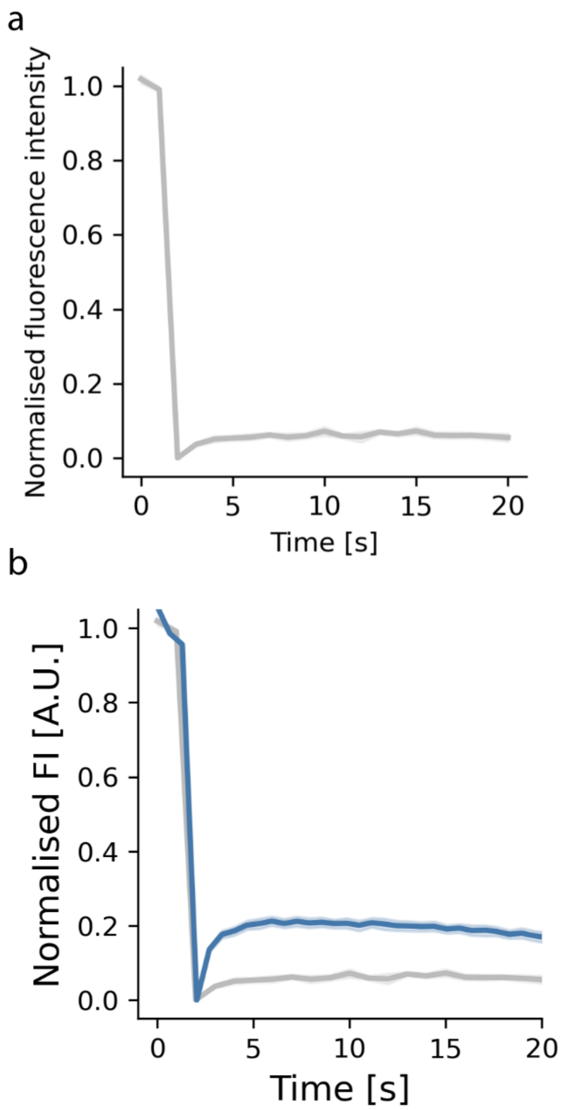


**Supplementary Fig S34: FRAP recovery curves of HERD-2.2-GFP droplets in cells under different growth conditions. a,** Fluorescence recovery of bleached spots within HERD-2.2–GFP protein condensates in *E. coli* grown at 18 °C after induction. Shaded area represents the standard error. n = 15 **b,** Overlay of fluorescence recovery of HERD-2.2–GFP droplets within *E. coli* grown at 37 °C after induction and chilled briefly to induce condensation (blue, n = 13), and at 18 °C after induction (grey).

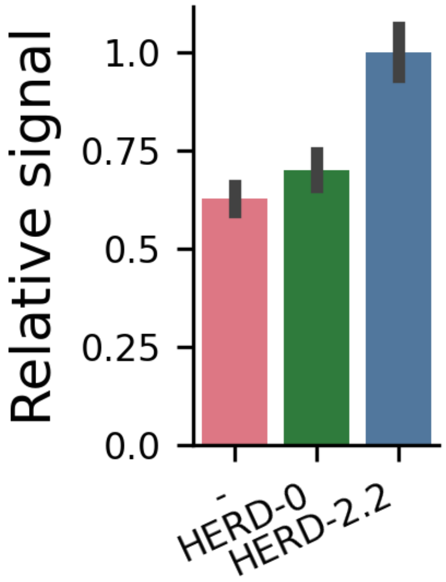


**Supplementary Fig S35: Quantification of expression levels of TnaA and FMO fusions by western blotting.** $OD_{700}$ adjusted cell pellets were collected at the same time as samples were collected for indigo measurement and used for western blotting to quantitatively compare the expression of the differentially tagged enzymes. Enzymes were western blotted against the His epitope tag from triplicate cultures and quantification performed in Image Studio Lite to measure enzyme expression. Background was set manually to avoid interference with the co-expressed GFP labelled polypeptides.

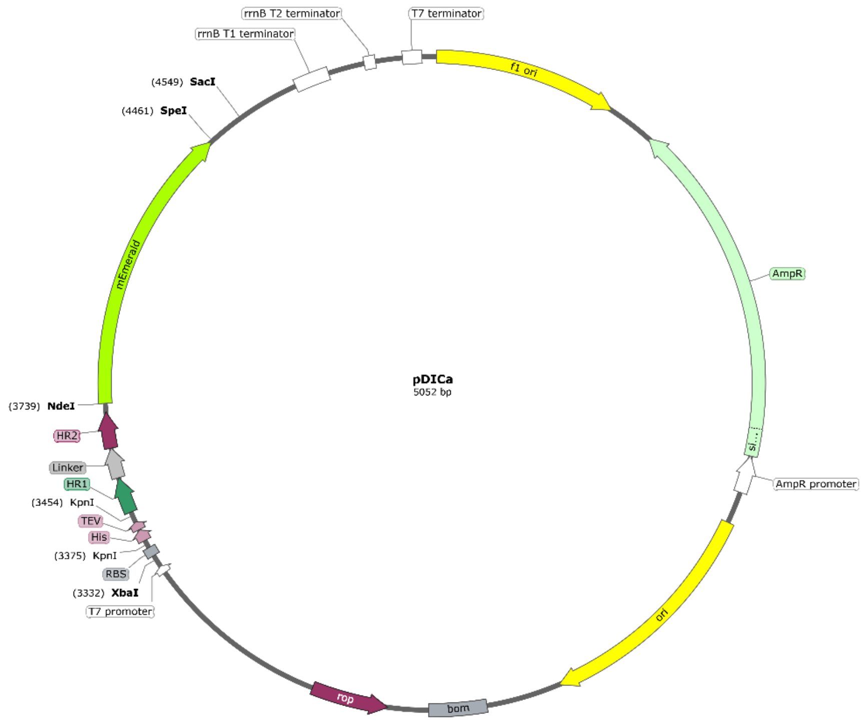


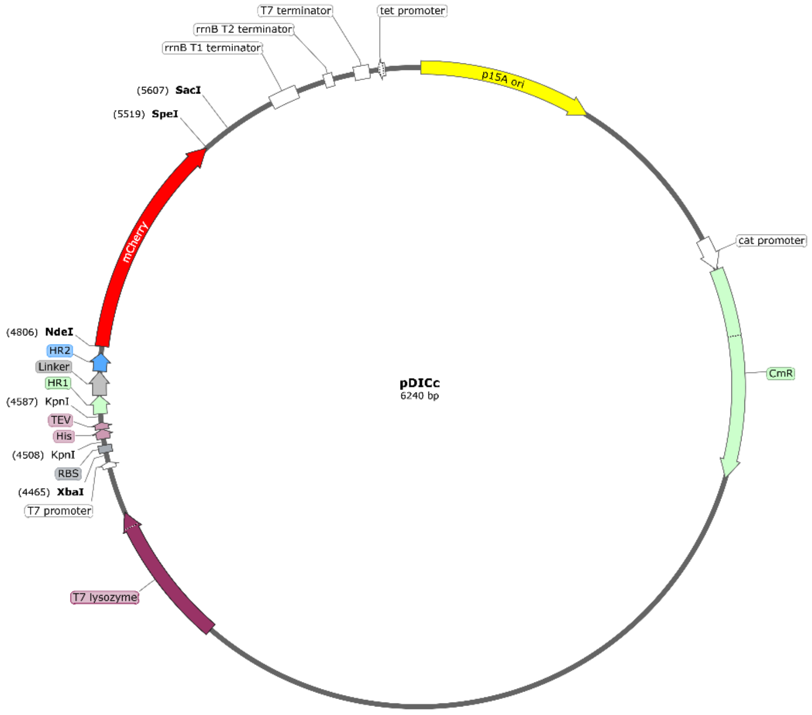


**Supplementary Fig S36: pDIC (design in cells) vectors used for protein expression.** Upper – pDICa (ampicillin selection). Lower – pDICc (chloramphenicol selection).
